# Ileitis abolishes tolerogenic functions of the enterohepatic bile acid pool

**DOI:** 10.64898/2026.09.13.751256

**Authors:** Zhiping Xiao, Koichi Sudo, Amber Delmas-Eliason, Jiabao Liu, Paola Munoz-Tello, Henry D. Dionne, Fiona Nguyen, Alexander D. Hondros, Huijuan Yang, B. JoNell Hamilton, Akshaya Balasubramanian, Courtney L. Hegner, Gillian Jacobsen, Erin M. Tonzi, Ila K. Kaul, Shannon Soucy, Corey A. Siegel, George A. O’Toole, Daniel Schultz, Casey T. Weaver, Alex Rodriguez-Palacios, Fabio Cominelli, Henry M. Krause, Maria T. Abreu, Paul A. Dawson, Douglas J. Kojetin, Mark S. Sundrud

## Abstract

Bile acids (BAs) regulate lipid uptake, epithelial integrity, and immune responses in the gut. Hepatocytes synthesize primary BAs, which microbiota metabolize into secondary metabolites. Together, these species form a composite pool that circulates enterohepatically between the liver and intestines. Here, we show that immuno-regulatory outputs of the enterohepatic BA pool involve competition between multiple BA species for individual nuclear receptors and are dependent on intestinal health. In healthy mice, the primary BA, tauro-β-muricholic acid (tβMCA) antagonized RORγt-mediated Th17 function in the presence of two secondary BAs (tDCA, tLCA) which activated RORγt. Conversely, ileitis in Crohn’s disease patients and *Tnf*^ΔARE/+^ mice depleted primary BAs by reducing the number and function of BA-transporting enterocytes. In mice, ileitis-driven tβMCA depletion enhanced tDCA/tLCA-mediated RORγt activation and supported Th17 cell function, whereas replenishing tβMCA in *Tnf*^ΔARE/+^ mice reestablished BA regulation of ileal Th17 cells. Thus, intra-pool competition underpins BA immunoregulatory functions and may reveal new opportunities for precision Crohn’s disease therapy.

## Introduction

Bile acids (BAs) have emerged as direct immunoregulatory metabolites in the gastrointestinal (GI) tract. All BAs derive from cholesterol catabolism in hepatocytes—and thus share a steroid backbone hydroxylated at the carbon-3 (C-3) position—whereas individual BA structures differ with regard to presence, absence or stereochemistry of other modifications. Nuclear receptors (NRs) are a common class of host BA receptors, including in immune cells. It has been posited that BA structural modifications may have co-evolved with host receptors, in part, to facilitate crosstalk between hepatocytes, enterocytes, gut microbiota, and intestinal immune cells.

Studies of BA-dependent intestinal immune regulation have centered on roles of single BA species. However, it is rarely considered how BA concentrations used in functional studies relate to physiological levels in gut mucosal tissues, which differ from those in gut luminal segments given limiting rates of active and passive absorption in the ileum, and of only passive absorption in the colon.^1^ Further, simplified models of single BA-NR interactions occurring in isolation are incongruent with physiological reality, where intestinal immune cells *in situ* are exposed to complex and dynamically regulated BA pools; BA species that circulate together in the same enterohepatic pool can bind the same NR but elicit opposing functions (*e.g.*, FXR is activated by tCDCA but inhibited by tβMCA^2,3^). Thus, physiological functions of BAs, like those of microbiota, reflect collective outputs of complex pools.

We recently described an approach to empirically measure the pool of endogenous BAs that circulates enterohepatically through the ileal mucosa of mice.^4^ In 30-week-old C57BL/6J (B6) wild type mice, this pool contains at least 42 BA species, and its total concentration in the fasted state totals more than 300 μM.^4^ Here, we show that spontaneous inflammation of the ileum (*i.e*., ileitis)—a common and intractable feature of Crohn’s disease (CD) modelled experimentally in *Tnf^ΔARE/^*^+^ mice, which overexpress TNFα due to ablation of a destabilizing 69-bp A/U-rich element (ARE) in the *Tnf* 3’ UTR^5^—reshapes ileal BA absorption dynamics to produce a smaller and more hydrophobic enterohepatic BA pool. By reconstituting physiologically defined ‘healthy’ and ‘ileitis-associated’ mouse enterohepatic BA pools from a library of commercially synthesized species, we identify three BAs that operate within these pools to directly, differentially and competitively modulate Th17 cell function via RORγt. One of these BAs, tβMCA, is a low-affinity RORγt antagonist synthesized in hepatocytes and maintained at high enterohepatic levels due to efficient absorption by ileal enterocytes in wild type, but not *Tnf^ΔARE/^*^+^, mice. The other two species—tDCA and tLCA—are higher affinity, but also less abundant, RORγt agonists produced, in part, via gut microbial metabolism whose ileal absorption remains more intact during ileitis. This integrated view of the physiological interplay between BA enterohepatic circulation and intestinal immune regulation establishes intra-pool BA-receptor competition as a defining feature of BA physiological functions and offers new insights into the pathogenesis and treatment of CD.

## Results

### Ileitis remodels the enterohepatic BA pool

Specialized enterocytes in the terminal ileum express the BA reuptake transporter, ASBT/*SLC10A2*, and maintain a large pool of enterohepatically cycling BAs.^6,7^ At steady-state, ∼60% of this pool consists of ‘primary’ (1°) BAs [*e.g.*, cholic acid (CA) and chenodeoxycholic acid (CDCA) in humans; CA, CDCA and alpha- or beta-muricholic acid (α/βMCA) in mice] that are side chain-conjugated to amino acids—glycine or taurine (*e.g*., gCA or tCA)—and that hepatocytes synthesize and secrete into the proximal small intestine (SI; duodenum) to enable lipid absorption. The remainder of the enterohepatic pool consists mostly of ‘secondary’ (2°) BA species and metabolites produced via microbial metabolism of 1° BAs in the large intestine (LI; cecum, colon) and that become incorporated into the enterohepatic pool following passive absorption in the LI (colonocytes lack ASBT expression) and re-secretion by the liver.

Given the unique function of the ileum in maintaining a distinctive pool of enterohepatically cycling BAs,^7^ we sought to define how ileitis affects the size and complexity of this pool. Using the same approach as reported in wild type mice,^4^ we used targeted mass spectrometry to quantify absolute concentrations of 88 BA species and metabolites (**Figure S1A**) in peripheral blood (pB) plus three GI sites—SI lumen content (siLC), superior mesenteric vein blood (smvB), feces—of *Tnf^ΔARE/^*^+^ mice with ileitis. *Tnf^ΔARE/^*^+^ mice were weaned and co-housed together with wild type littermates to normalize microflora; BAs were measured at 30-weeks of age, when 100% of *Tnf^ΔARE/^*^+^ mice display severe ileitis (see below). Consistent with impaired intestinal BA absorption, total BA levels in *Tnf^ΔARE/^*^+^ *vs*. wild type mice were ∼3.5x lower in both siLC (which reflects BAs made and/or secreted by hepatocytes into the SI) and smvB (which captures BAs absorbed through the ileal mucosa and recirculating to the liver) (**Figure 1A**). Conversely, total BA levels in non-enterohepatic tissues (*i.e.,* feces, pB) were unaffected by ileitis in *Tnf^ΔARE/^*^+^ mice (**Figure 1A**). Within the smvB (enterohepatically cycling) BA pool, ileitis in *Tnf^ΔARE/^*^+^ mice depleted 1° and 2° conjugated (c)BAs, whereas 2° unconjugated (u)BA species and metabolites were better retained and became proportionately enriched (**Figure 1B**). These shifts produced a more diverse and hydrophobic enterohepatic BA pool in *Tnf^ΔARE/^*^+^ mice, with features more akin to normal colonic BA pools that passively transit the colonic mucosa and recirculate to the liver via the inferior mesenteric vein (imv) (**Figure 1C-D**).

**Figure 1.**
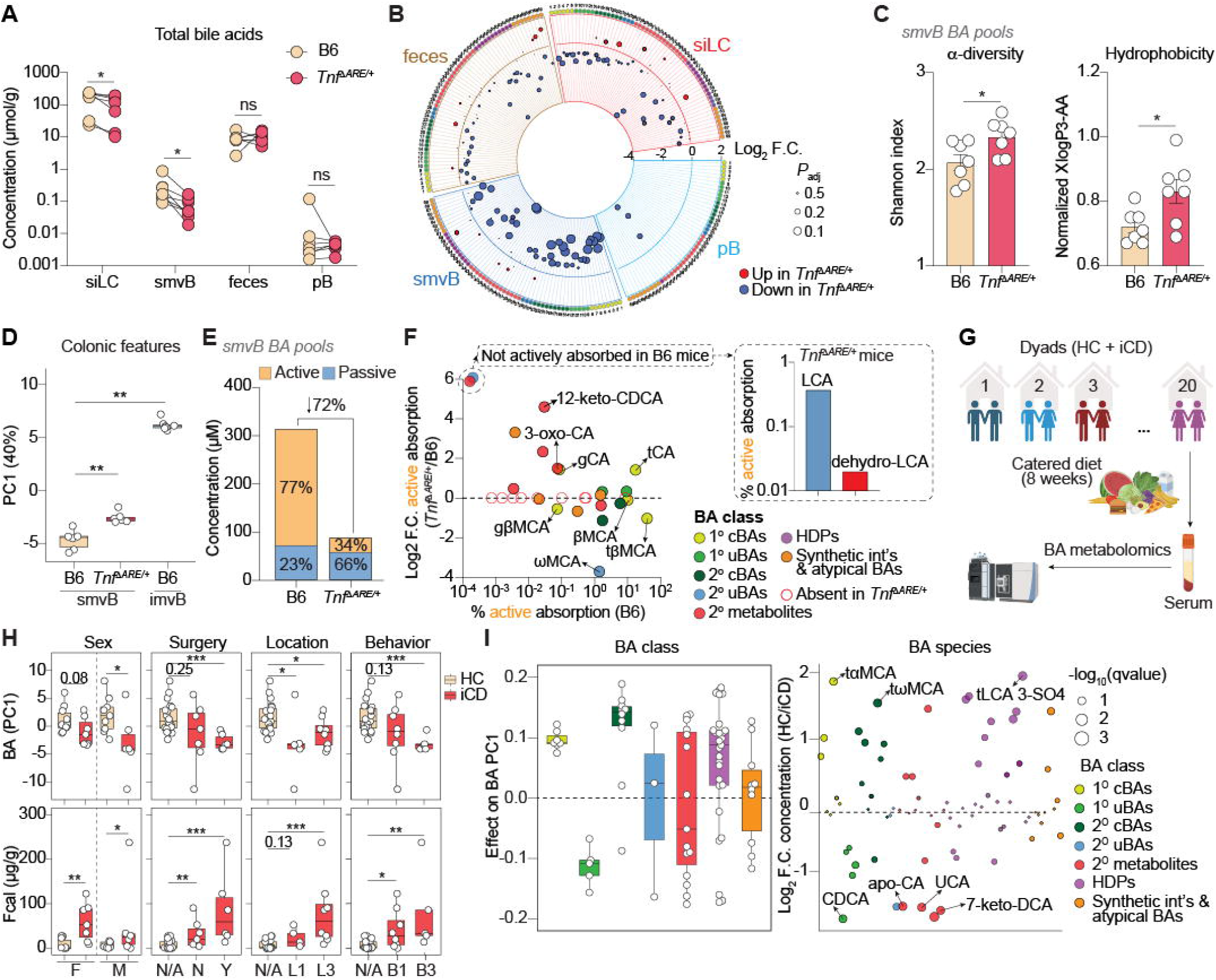
Ileitis reshapes BA enterohepatic circulation. (A) Total BA concentrations in 4 sites from 7 pairs of co-housed C57BL/6 wild type (B6) or *Tnf^DARE/^*^+^ mice. siLC, small intestinal lumen content; smvB, superior mesenteric vein blood; pB, peripheral blood. (B) Log_2_ fold-changes (Log_2_ F.C.) in mean BA concentrations (*n* = 7 mice) from the 4 sites in (A) in *Tnf^DARE/^*^+^ *vs.* B6 mice. Only BAs with median concentrations >0 in each site are shown; species are numbered and color-coded by class as in **Figure S1A**. Red or blue dots indicate species increased or decreased in *Tnf^DARE/^*^+^ *vs.* B6 sites, respectively. (C) α-diversity and normalized hydrophobicity of smvB BA pools from B6 or *Tnf^DARE/^*^+^ mice (*n* = 7 mice). α-diversity (*left*) quantified by Shannon index; hydrophobicity (*right*) was calculated by dividing cumulative XlogP3-AA values for each pool by total concentration. (D) Principal Component Analysis (PCA) of BA pools measured in smvB or predicted in inferior mesenteric vein blood (imvB) of B6 or *Tnf^DARE/^*^+^ mice (*n* = 6-7 mice). (E) Average amounts of active (tan) and passive (blue) ileal BA absorption in B6 or *Tnf^DARE/^*^+^ mice. Numbers indicate percentages of total ileal BA absorption; active ileal absorption is reduced by 72% in *Tnf^DARE/^*^+^ mice. (F) Log_2_ F.C. in active ileal BA absorption (*n* = 7 mice), per species, in *Tnf^DARE/^*^+^ *vs*. B6 mice (y-axis) relative to their percentage of total active ileal BA absorption in B6 mice (x-axis). Species are color-coded by class as in (B) and **Figure S1A**. Empty red circles show species that are actively absorbed in only B6 mice; two species (insert at right; LCA, dehydro-LCA) are only actively absorbed in *Tnf^DARE/^*^+^ mice. 1° cBAs, primary conjugated BAs; 1° uBAs, primary unconjugated BAs; 2° cBAs, secondary conjugated BAs; 2° uBAs, secondary unconjugated BAs; HDPs, hepatic phase 2 detoxification products. All BA measurements were performed in 30-weeks old B6 or *Tnf^DARE/^*^+^ mice co-housed at weaning. (G) Serum BA profiling was performed on co-habitating human healthy controls (HC) or ileal Crohn’s disease (iCD) patients fed an identical (catered) diet for 8 weeks. (H) Serum BA principal component 1 (PC1) values (*top*, as in **Figure S2C**) and fecal calprotectin (*fcal*) levels *bottom*) in iCD patients *vs*. HC broken out by biological sex (F, female; M, male); history of ileal resection (surgery); Montreal classification of CD location (L1, ileal; L3, ileocolonic); Montreal classification of CD behavior (B1, inflammatory, not stricturing or penetrating; B3, penetrating). (I) *Left*, contributions of BA classes [abbreviated and color-coded as in (B and F) and **Figure S1A**] to BA PC1 values (eigenvectors) in HC and iCD patients. *Right*, Log_2_ F.C. in mean serum concentrations of individual BA species (*n* = 17; color-coded by BA class as above) from iCD patients *vs*. HC. Hydrophobicity was calculated using XLOGP3-AA values from PubChem. Data are presented as the mean ± SEM. Statistical significance was determined by Wilcoxon matched-pairs signed rank test (A), paired *t*-test (C), and Wilcoxon signed-rank test (B, D, H, I). \**P* < 0.05, \*\**P* < 0.01, \*\*\**P* < 0.001, ns, not significant. See also **Figure S1** and **Figure S2**.

We previously showed that quantitative relationships between siLC and smvB BA levels within individual wild type mice reflect ileal absorption rates; by defining these rates for each BA species, and in mice that have or lack Asbt/*Slc10a2*, contributions of active (Asbt-mediated) BA absorption *vs*. passive diffusion can be parsed and quantified *in vivo* (**Figure S1B**).^4^ To further examine the mechanisms by which ileitis perturbs ileal BA absorption, we crossed *Tnf^ΔARE/^*^+^ mice with Asbt-deficient (*Slc10a2*^-/-^) animals and compared rates and routes of ileal BA absorption in *Tnf^ΔARE/^*^+^ *vs*. wild type mice.^4^ Ileitis in *Tnf^ΔARE/^*^+^ mice reduced the total amount of active (Asbt-mediated) BA absorption, whereas passive absorption (*i.e*., diffusion) was unchanged (**Figure 1E**). More interestingly, ileitis also altered which BA species were absorbed by Asbt in the ileum, with a shift away from conjugated and unconjugated muricholic acids (α/β/ωMCAs) and towards conjugated cholic acids (tCA, gCA), 2° uBAs and 2° metabolites (**Figure 1F**). Nine BA species were only absorbed by Asbt in healthy wild type mice, and two other species—lithocholic acid (LCA) and the related bacterial metabolite, dehydro-LCA—were only absorbed by Asbt during ileitis (**Figure 1F**). These results indicate that ileitis fundamentally alters Asbt transport dynamics to remodel the enterohepatic BA pool.

Enrichment of microbially generated 2° uBAs and metabolites within the enterohepatic pool of inflamed *Tnf^ΔARE/^*^+^ mice suggested that dysbiosis in this model may have limited impacts on microbiota-mediated BA metabolism. In line with this, few community- or species-level differences were observed in the abundance of key BA-metabolizing genes (*Bsh*, *7ahsdh*) within cecal microbiota of co-housed *Tnf^ΔARE/^*^+^ and wild type mice (**Figure S1C-D**). Unencumbered bacterial BA metabolism in *Tnf^ΔARE/^*^+^ mice with ileitis is consistent with the lack of severe colonic inflammation, where most bacterial BA metabolism occurs,^5,8^ and is distinct from mouse colitis models where dysbiosis is consistently associated with reduced synthesis of 2° uBA species and metabolites.^9–11^

Notable differences exist between BA metabolic pathways in humans and mice, whereas the process of enterohepatic circulation—mediated by conserved ileal and liver BA transporters—is more similar.^12–14^ To understand if ileitis-associated shifts in BA enterohepatic circulation in *Tnf^ΔARE/^*^+^ mice reflects pathophysiology of ileal (i)CD in humans, we analyzed the same panel of 88 BAs in serum collected from 17 pairs (dyads) of adult healthy controls (HCs) and iCD patients living in the same household and fed an identical (catered) diet for 8 weeks (**Figure 1G**).^15^ We focused on serum for this study, as the pool of enterohepatically cycling BAs in smvB is not accessible in humans outside of complex GI surgeries and is not reflected by BAs in stool (**Figure 1A)**. Further, as diet and microbiota are the two main environmental sources of BA variability in humans—and microbiota tend to be shared within households^16^—this study design afforded a more controlled and rigorous analysis of iCD-specific serum BA features.

Total serum BA levels trended, but were not significantly, lower in iCD patients *vs*. housemate HCs (**Figure S2 A-B**). This was not unexpected, as: (*i*) we observed few gross differences in pB BAs between co-housed wild type and *Tnf^ΔARE/^*^+^ mice (**Figure 1A**); and (*ii*) we previously showed that BAs in mouse pB reflect a mixture of species that escape first-pass hepatic reuptake after absorption in either the ileum or colon.^4^ Thus, we next used principal component analysis (PCA) to scan for serum BA sub-signatures that more directly reflect the pool of human BAs absorbed through the ileum (**Figure S2C**). PC1 explained the most variance across all samples (20.8%), was significantly different between iCD patients and healthy controls, and—similar to an established protein biomarker of IBD, fecal calprotectin (Fcal)^17^—consistently discriminated iCD patients from HCs regardless of biological sex, prior intestinal surgery or whether disease was simple/inflammatory (Montreal class B1) or complex/penetrating (B3) (**Figure 1H**).^18^ Interestingly, BA PC1 preferentially identified CD patients with ileal-only disease (Montreal location L1), whereas increased Fcal, as expected, was better at discriminating CD with colonic involvement (L3) (**Figure 1H**). At BA class and species levels, 1° and 2° cBAs—major ASBT transport substrates reduced in enterohepatic circulation of *Tnf^ΔARE/^*^+^ mice—contributed strongly to PC1 and were reduced in iCD *vs*. HC sera (**Figure 1B, 1I**). Conversely, 1° uBAs and 2° metabolites—direct products of gut microbial BA metabolism increased in enterohepatic circulation of *Tnf^ΔARE/^*^+^ mice—contributed oppositely to PC1 and were elevated in iCD patients (**Figure 1B, 1I**). These results suggest that aberrant ileal BA absorption is a core biological feature of ileitis and could inform development of iCD-specific serum biomarkers.

### Ileitis depletes and reprograms ASBT-expressing ileocytes

Consistent with the marked and specific impacts of ileitis on active (Asbt-mediated) BA absorption in *Tnf^ΔARE/^*^+^ mice (**Figure 1E** and **1F**), Asbt/*Slc10a2* gene and protein expression were reduced in ileal tissue of *Tnf^ΔARE/^*^+^ *vs*. wild type mice, and the extent of Asbt/*Slc10a2* downregulation correlated with increasing *Tnf* expression (**Figure 2A** and **Figure S3A**). Likewise in humans, ASBT/*SLC10A2* gene expression was reduced in a large and independent set of iCD *vs*. UC patient ileal biopsies (UC does not involve the ileum so served as ‘controls’ here), with lowest ASBT/*SLC10A2* expression again observed in the most inflamed biopsies (**Figure 2B**).

**Figure 2.**
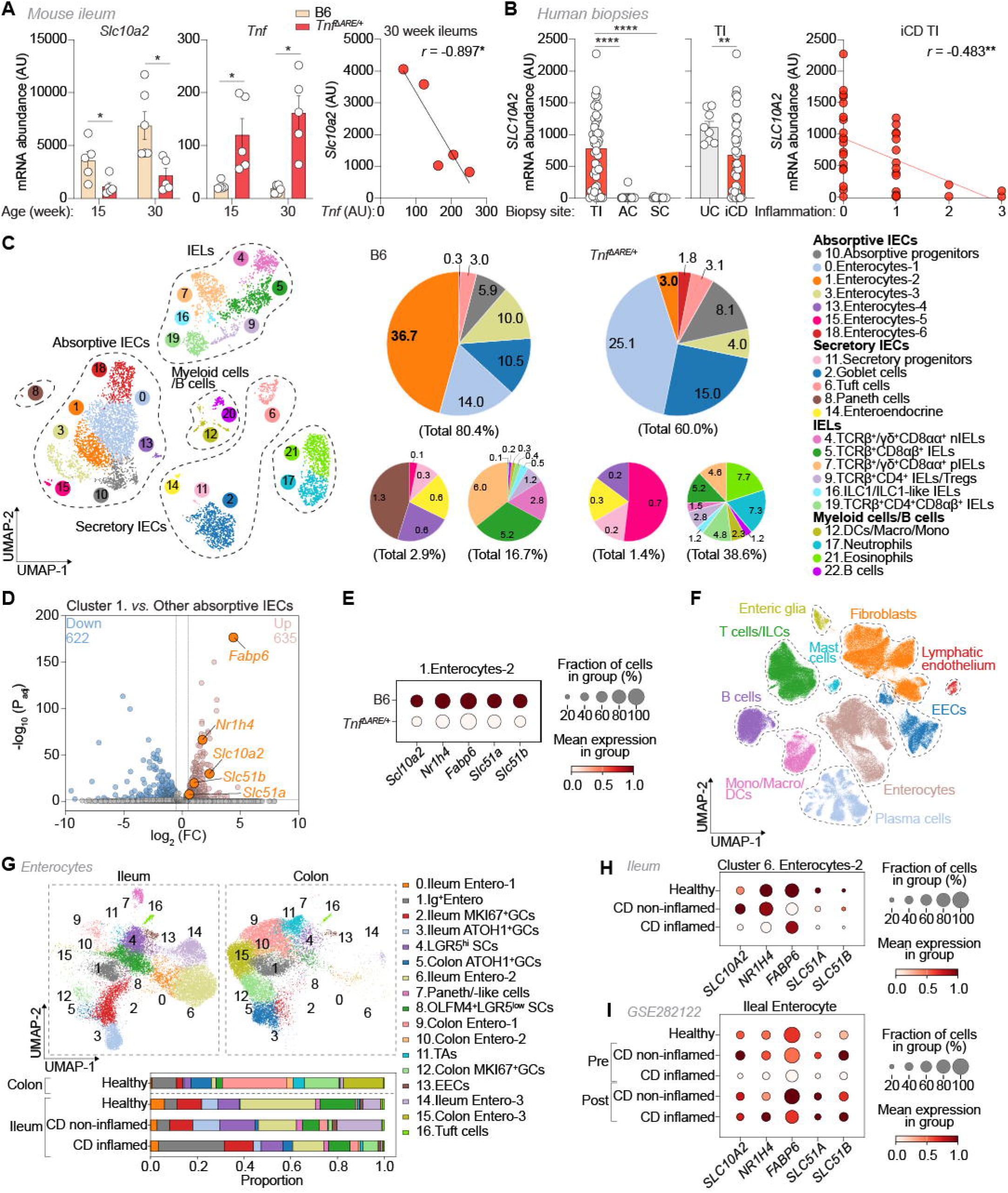
ASBT*/SLC10A2*-expressing enterocytes are transcriptionally reprogrammed in ileitis. (A) mRNA expression of *Slc10a2* and *Tnf* in terminal ileum from B6 and *Tnf^ΔARE/^*^+^ mice. B6 and *Tnf^ΔARE/^*^+^ mice were co-housed at weaning and analyzed at 15-weeks and 30-weeks of age (*n* = 5-6 mice). *Slc10a2* (*left*), *Tnf* (*middle*), correction analysis between *Slc10a2* and *Tnf* expression (*right*). (B) mRNA expression of *SLC10A2* in biopsies from patients with ileal Crohn’s disease (iCD) or Ulcerative colitis (UC). *SLC10A2* expression at different location (*left*, TI, terminal ileum, *n* = 45; AC, ascending colon, *n* = 27; SC, sigmoid colon, *n* = 14), *SLC10A2* expression in terminal ileum from iCD and UC (*middle*), correlation of *SLC10A2* expression and inflammatory scores (*right*). (C) scRNA-Seq of ileal epithelial fractions from B6 and *Tnf^ΔARE/^*^+^ mice. Uniform manifold approximation and projection (UMAP, *left*), percentages of each cell cluster in B6 and *Tnf^ΔARE/^*^+^ mice (*right*). IECs, intestinal epithelial cells; nIELs, natural intraepithelial lymphocytes; pIELs, peripheral intraepithelial lymphocytes; ILC, Innate lymphoid cells; DCs, Dendritic cells; Macro, Macrophages; Mono, Monocytes. (D) Volcano plot of differentially expressed genes (DEGs) between Cluster 1 and Other absorptive IECs. Pink (*upregulated*) and blue (*downregulated*) dots denote significant differences with *P*_adj_ < 0.05 and log_2_ (FC) > 0.50. BA-metabolizing genes are highlighted in orange. (E) Dot plot of BA-metabolizing genes in Cluster 1 from B6 and *Tnf^ΔARE/^*^+^ mice. (F) UMAP of enterocytes and lamina propria cells from the ileum and colon of healthy controls (Healthy) and Crohn’s diseases (CD) patients. (G) Subclustering of enterocytes and percentage of enterocytes subsets in ileum and colon from healthy controls and CD patients. Ileal enterocytes (*top left*), colonic enterocytes (*top right*), percentage of enterocytes subsets in Colon (Healthy) and ileum (Healthy, CD non-inflamed, and CD inflamed) (*bottom*). (H) Dot plot of BA metabolizing genes in ileal Cluster 6 enterocytes (Enterocytes-2) from Health, non-inflamed, or inflamed CD patients. (I) Dot plot of BA-metabolizing genes in ileal enterocytes from IBD patients before (pre) and after (post) treatment with adalimumab.^20^ scRNA-Seq for ileal epithelial fractions was performed in mice at 15 weeks of age with two mice pooled per sample. Data are presented as the mean ± SEM. Statistical significance was determined by two-tailed unpaired Welch’s *t*-test (A and B, *middle*), one-way ANOVA with Tukey’s post hoc test (B, *left*), and Wilcoxon signed-rank test (D-E and G-I). \**P* < 0.05, \*\**P* < 0.01, \*\*\**P* < 0.001, \*\*\*\**P* < 0.0001. See also **Figure S3** and **Figure S4**.

Reduced ASBT expression and function in inflamed ileums of humans and mice could result from increased turnover (death) of mature ASBT-expressing enterocytes and/or transcriptional reprogramming of viable cells. To examine both possibilities, we performed parallel single cell (sc)RNA-seq analyses of ileal epithelial cells from mice (wild type, *Tnf^ΔARE/^*^+^) or humans (HC, iCD) with or without ileitis (**Figure S3B**). In mice, seven clusters of absorptive enterocytes were identified and annotated based on known marker genes, in addition to secretory epithelial (goblet, Paneth) cells and a variety of epithelium-associated immune cell lineages (**Figure 2C** and **Figure S3C**). Sub-clustering of absorptive enterocytes revealed clusters 0, 1 and 2 expressed varying levels of Asbt/*Slc10a2* (**Figure S3D**). Cluster 1 was the most abundant enterocyte cluster at steady-state and expressed the highest levels of Asbt/*Slc10a2*, as well as other hallmarks of BA-absorptive ileocytes (*e.g*., FXR/*Nr1h4*, *Fabp6*, *Slc51a*, *Slc51b*) (**Figure 2C-D, Figure S3E**-**S3G**). Fewer cells from *Tnf^ΔARE/^*^+^ mice contributed to cluster 1, compared with wild type counterparts and consistent with inflammation-induced turnover (**Figure 2C**). At the same time, viable cluster 1 cells from *Tnf^ΔARE/^*^+^ mice displayed lower expression of Asbt/*Slc10a2*, FXR/*Nr1h4*, *Fabp6*, *Slc51a* and *Slc51b*, suggesting inflammation-associated transcriptional reprogramming (**Figure 2E**). Similar results were observed in humans; the largest cluster of ileal enterocytes (cluster 6 in **Figure 2F-G**) showed distinctively high expression of BA-related genes, including ASBT/*SLC10A2*, was not present in colonic biopsies, was proportionately reduced in iCD *vs*. HC ileal biopsies, and displayed ileitis-associated reductions in ASBT/*SLC10A2* expression (**Figure 2G-H, Figure S4**).

Downregulation of both ASBT/*SLC10A2* and FXR/*NR4H1* within absorptive enterocytes from inflamed human and mouse ileums was notable because, at steady-state, BA activation of FXR in ileal enterocytes represses ASBT/*SLC10A2* expression to decrease the levels of BAs in enterohepatic circulation.^19^ Thus, we postulated that inflammatory cytokines produced during ileitis might contribute to ASBT/*SLC10A2* downregulation. Indeed, by analyzing a published scRNA-seq dataset^20^, we found that reduced enterocyte ASBT/*SLC10A2* expression in inflamed CD patient ileal biopsies was increased following anti-TNFα biologic therapy, even in tissues that remained histologically inflamed (**Figure 2I**). Thus, a combination of enterocyte turnover and cytokine-mediated transcriptional reprogramming may precipitate formation of aberrant enterohepatic BA pools during ileitis.

### Enterohepatically circulating BAs in *Tnf^ΔARE/^*^+^ mice fail to repress Th17-associated ileitis

Changes in the size or composition of enterohepatically cycling BA pools are likely to change immunoregulatory networks within the ileal mucosa.^21^ To determine if the aberrant pool of enterohepatically cycling BAs in *Tnf^ΔARE/^*^+^ mice regulates ileitis pathogenesis, we next blocked residual ileal BA absorption in *Tnf^ΔARE/^*^+^ mice using both pharmacologic and genetic approaches. Treatment of 25-week-old (already-inflamed) *Tnf^ΔARE/^*^+^ mice with the BA sequestering resin, cholestyramine (CME),^22^ reduced ileal histopathology after 5 weeks (**Figure S5A-B**). Consistent with this, *Tnf^ΔARE/^*^+^ mice bred to lack Asbt-dependent BA absorption lived 2- to 3-months longer than Asbt-sufficient cage-mates and showed delayed onset of ileitis (**Figure 3A-B, Figure S5C**). Elevated *Tnf* expression, a direct consequence of the *Tnf^ΔARE/^*^+^ mutation owing to increased half-life of *Tnf* mRNAs^5^, was similarly high in whole ileal tissue of both Asbt-sufficient and Asbt-deficient *Tnf^ΔARE/^*^+^ mice (**Figure 3C**), suggesting that enterohepatically cycling BAs regulate functions of inflammatory cells in the ileum beyond TNFα production. Cecal microbiota in co-housed Asbt-sufficient and Asbt-deficient *Tnf^ΔARE/^*^+^ mice were also similar, as determined by shotgun metagenomics sequencing, and both genotypes were equivalently colonized with the ileal Th17 cell-induced commensal bacteria, segmented filamentous bacteria (SFB) (**Figure S5D-E**).^23^

**Figure 3.**
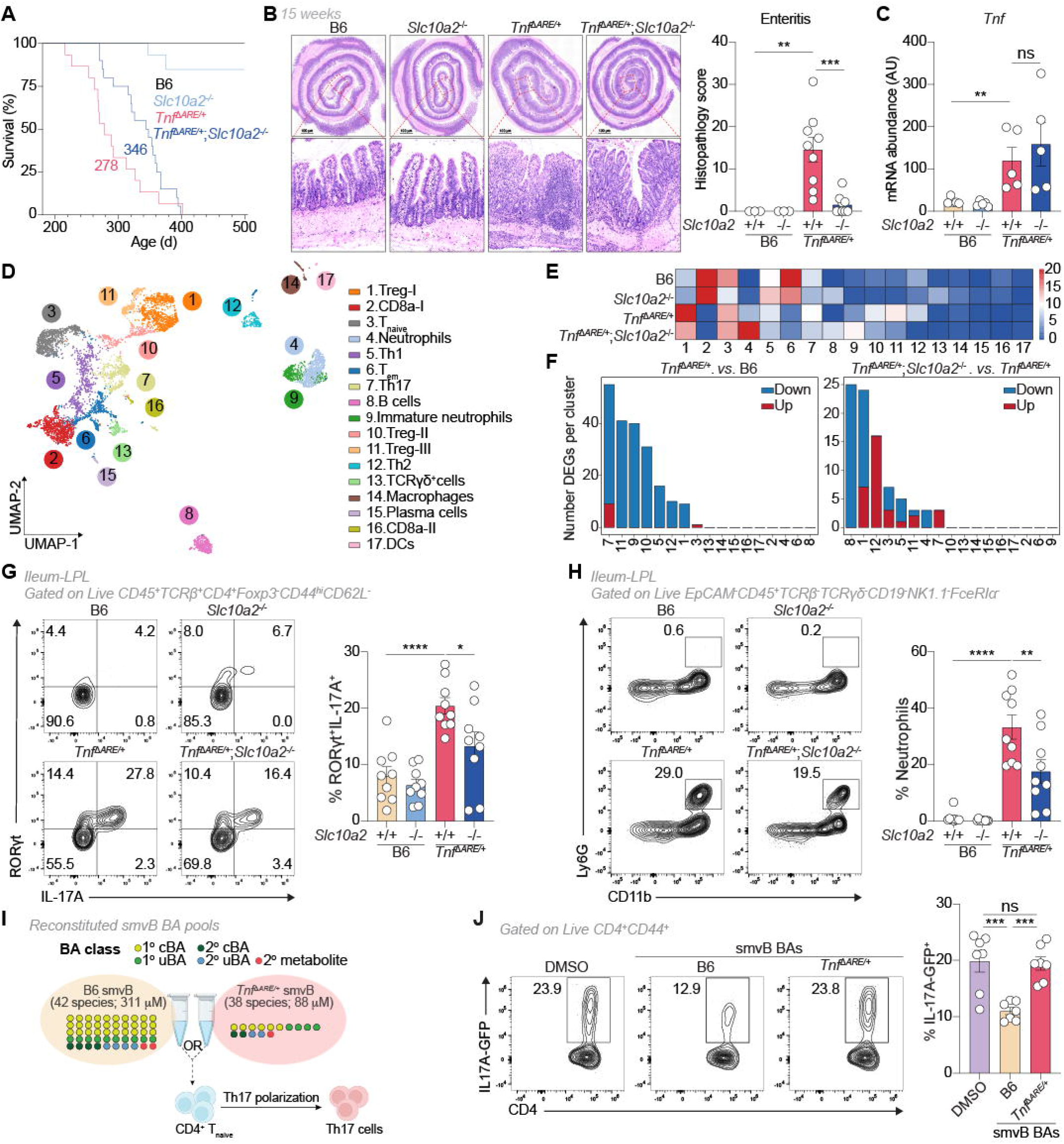
Bile acid pools support, rather than suppress, ileal Th17 cells during ileitis. (A) Survival curve of B6 (n = 13 mice), *Slc10a2*^-/-^ (*n* = 15 mice), *Tnf^ΔARE/^*^+^ (*n* = 15 mice), and *Tnf^ΔARE/^*^+^;*Slc10a2*^-/-^ (*n* = 20 mice) mice. Mice were co-housed at weaning and monitored up to 500 days. (B) Representative hematoxylin and eosin (H&E) staining of terminal ileum. B6 (*n* = 3), *Slc10a2*^-/-^ (*n* = 3), *Tnf^ΔARE/^*^+^ (*n* = 11), and *Tnf^ΔARE/^*^+^;*Slc10a2*^-/-^ (*n* = 13 mice) mice were co-housed at weaning and analyzed at 15-weeks of age. Representative images (*left*, Scale bar, 100 μm), quantification of inflammatory scores (*right*). (C) mRNA expression of *Tnf* in terminal ileum. (D) scRNA-Seq UMAP of immune cells from ileal lamina propria. B6, *Slc10a2*^-/-^, *Tnf^ΔARE/^*^+^, and *Tnf^ΔARE/^*^+^;*Slc10a2*^-/-^ mice were co-housed at weaning. TCRβ^+^CD4^+^ T cells and CD4^-^ cells were sorted and mixed with 1:1 ratio for scRNA-Seq. Two mice pooled per sample. (E) Heatmap of percentage of each cell cluster. (F) Number of differentially expressed genes (DEG) in clusters. *Tnf^ΔARE/^*^+^. *vs*. B6 mice (*left*), *Tnf^ΔARE/^*^+^;*Slc10a2*^-/-^. *vs*. *Tnf^ΔARE/^*^+^ mice. *P_adj_* < 0.05. (G) Flow cytometric analyses of IL-17A-producing Th17 cells. RORγt^+^IL-17A^+^ cells were gated on Live CD45^+^TCRβ^+^CD4^+^CD44^hi^CD62L^-^Foxp3^-^ cells. Representative flow plots (*left*), frequency of RORγt^+^IL-17A^+^ cells (*right*). (H) Flow cytometric analyses of neutrophils. CD11b^+^Ly6G^+^ cells were gated on Live CD45^+^EpCAM^-^TCRβ^-^TCRγδ^-^CD19^-^FceRIa^-^NK1.1^-^ cells. Representative flow plots (*left*), frequency of CD11b^+^Ly6G^+^ cells (*right*). (I) Schematic of *in vitro* non-pathogenic Th17 (npTh17) polarization and treatment of smvB BA pools of B6 (B6 BAs) or *Tnf^ΔARE/^*^+^ (*Tnf^ΔARE/^*^+^ BAs) mice. (J) Flow cytometric analyses of IL-17A-GFP^+^ cells. CD4^+^CD25^-^CD44^-^ T cells were purified from IL-17A-GFP reporter mice, polarized, and treated with DMSO (vehicle) or synthetic smvB BA pools of B6 or *Tnf^ΔARE/^*^+^ for 3 days (*n* = 7). IL-17A-GFP^+^ cells were gated on live CD4^+^CD44^+^ T cells. Representative flow plots (*left*), frequency of IL-17A-GFP^+^ cells (*right*). B6, *Slc10a2*^-/-^, *Tnf^ΔARE/^*^+^, and *Tnf^ΔARE/^*^+^;*Slc10a2*^-/-^ mice were co-housed at weaning and analyzed at 15-weeks of age (*n* = 5, C; *n* = 9, G to I). Data are presented as the mean ± SEM. Statistical significance was determined by one-way ANOVA with Tukey’s post hoc test (B and C, G to I). \**P* < 0.05, \*\**P* < 0.01. See also **Figure S5**, **Figure S6**, and **Figure S7**.

To explore mechanisms, we used scRNA-seq (**Figure S6A**) and spectral flow cytometry analyses to survey ileal lamina propria immune landscapes in co-housed wild type or *Tnf^ΔARE/^*^+^ mice with or without Asbt. scRNA-seq analysis discriminated 17 immune cell clusters, which we annotated based on known marker genes (**Figure S6B-C**) and which included multiple subsets of CD4^+^ T regulatory (Treg) and effector (Teff) cells, as well as CD8^+^ T cells, B cells (including plasma cells), dendritic cells (DCs), macrophages and granulocytes (**Figure 3D**). Two clusters of RORγt-expressing CD4^+^ T cells [Th17 cells (cluster 7) and RORγt^+^ Treg cells (cluster 11)] were proportionally enriched in the *Tnf^ΔARE/^*^+^ *vs*. wild type ileum and were decreased in *Tnf^ΔARE/^*^+^ ileums lacking Asbt-dependent BA absorption (**Figure 3E**). These same clusters displayed the most differential gene expression between *Tnf^ΔARE/^*^+^ *vs*. wild type ileums and were among 8 clusters demonstrating differential gene expression in the *Tnf^ΔARE/^*^+^ ileum with or without Asbt (**Figure 3F**), suggestive of BA-dependent gene regulation.

Reduced levels of both total (RORγt^+^) and IL-17A^+^ Th17 cells, but not RORγt^+^ Treg cells, in Asbt-deficient *vs*. control *Tnf^ΔARE/^*^+^ ileums were also evident by flow cytometry (**Figure 3G, Figure S7A-S7B**). Neutrophils, which are recruited to inflamed tissues by IL-17A-secreting Th17 cells, were also decreased in ileums of *Tnf^ΔARE/^*^+^ mice lacking Asbt-dependent BA absorption (**Figure 3H, Figure S7C**). Conversely, loss of Asbt-mediated ileal BA absorption in *Tnf^ΔARE/^*^+^ mice had no impact on numbers or percentages of Th17 cells or neutrophils in the colon (**Figure 3F-H, Figure S7**), consistent with a local mechanism of BA-dependent Th17 cell regulation in the ileum that follows Asbt-mediated absorption. As Th17 cells are genetically associated with CD risk in humans and are regulated by a sterol- and BA-sensing NR, RORγt. ^24–26^, these results suggested that enterohepatic BA pools may impinge directly on ileal immune cell function and inflammation at the level of Th17 cells and RORγt. To examine this possibility, we reconstituted enterohepatic BA pools measured in the smvB of wild type or *Tnf^ΔARE/^*^+^ mice (**Figure 1A, Table S1**) using a library of commercially synthesized BA species and added these pools to naive CD4^+^ T cells stimulated *in vitro* under Th17-polarizing conditions. Indeed, the wild type, but not *Tnf^ΔARE/^*^+^, enterohepatic BA pool directly repressed development of IL-17A^+^ Th17 cells (**Figure 3I-J**). Together, these results reveal a direct tolerogenic function of the enterohepatic BA pool in wild type mice that is abolished by ileitis, though they do not exclude the potential of other (less direct) effects of aberrant enterohepatic BA pools in *Tnf^ΔARE/^*^+^ mice on ileal immune cell function and inflammation (*e.g*., through changes to microbiota).

### Competitive BA-binding to RORγt shapes Th17 cell regulation by the enterohepatic BA pool

In exploring mechanisms of direct Th17 cell regulation by physiologically defined BA pools, we found that *in vitro* inhibition of IL-17A^+^ Th17 cell development was a unique function of the enterohepatic (smvB) BA pool in wild type mice; BAs reconstituted to mimic BA pools measured in non-enterohepatic sites of the same animals (*i.e.*, feces, peripheral blood) did not repress IL-17A^+^ Th17 cell development (**Figure S8A, Table S1**). Mechanistically, the wild type enterohepatic BA pool repressed development of IL-17A^+^ Th17 cells in manners that were unrelated to cytotoxicity or impaired proliferation, and this pool inhibited *Il17a* gene expression in both ‘pathogenic’ and ‘non-pathogenic’ Th17 cell cultures^27^ without suppressing expression of other effector or regulatory cytokines (**Figure S8B-I**). Within Th17 cells, the wild type enterohepatic BA pool did not impair STAT3 phosphorylation or RORγt upregulation, indicating that initial events in Th17 cell lineage commitment were intact, and inhibitory effects of the enterohepatic BA pool on Th17 cell IL-17A expression became amplified over time after RORγt upregulation (**Figure S8J-L**). These results suggested that Th17 cell regulation by the wild type enterohepatic BA pool may involve direct modulation of RORγt activity. Consistent with this, reconstituted enterohepatic BA pools of wild type, but not *Tnf^ΔARE/^*^+^, mice repressed human (h)RORγ ligand-binding domain (LBD)-mediated transcriptional reporter activity in transfected CHO cells, whereas neither pool repressed the activity of the related NR, RORα, and both pools—despite marked differences in overall size and complexity—similarly activated the canonical BA receptor, FXR (**Figure 4A**).

**Figure 4.**
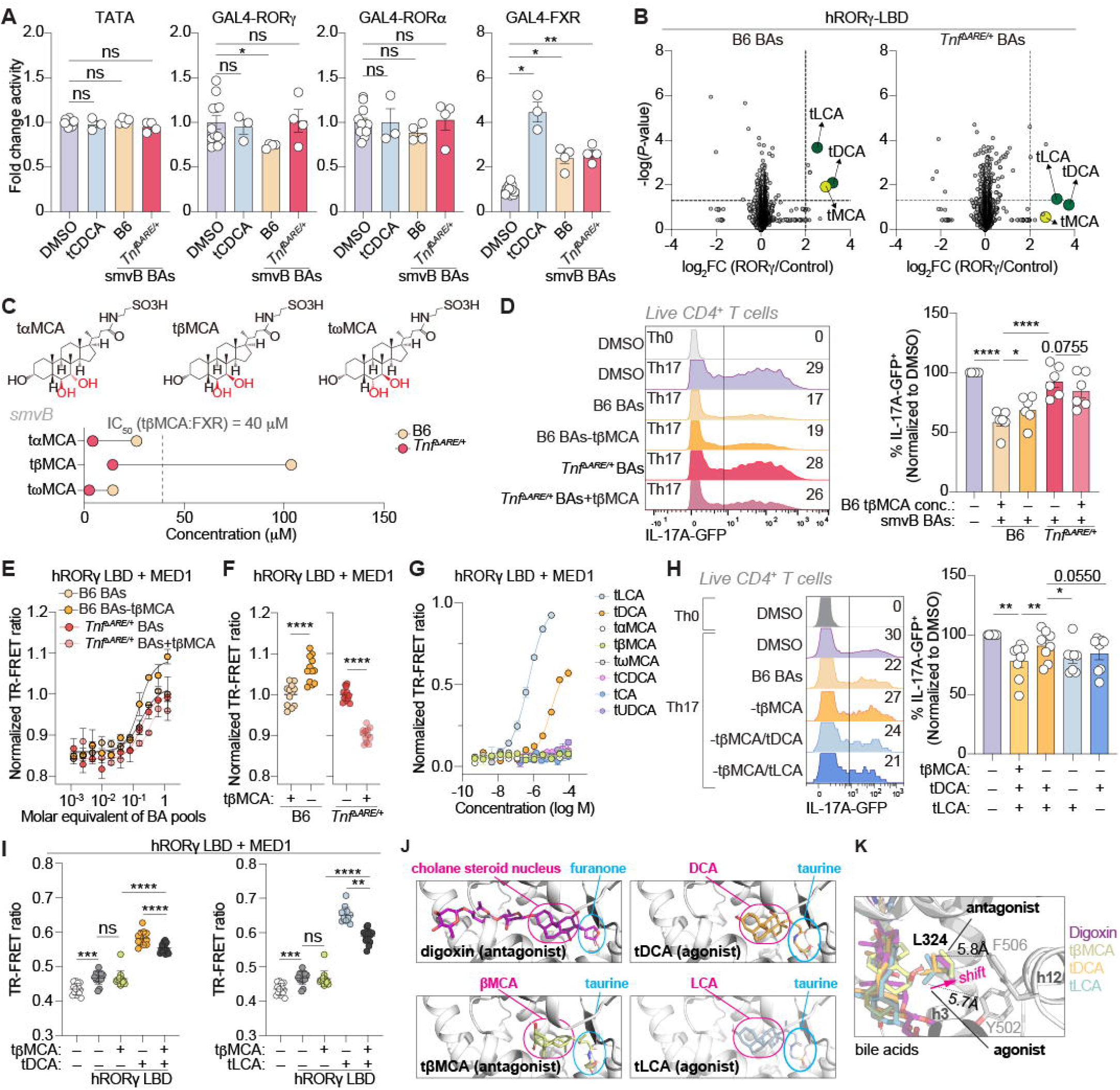
Enterohepatic BA pools directly and competitively regulate RORγ. (A) Activity of human nuclear receptors (NRs) with the treatments of tCDCA and BA pools. Chinese hamster ovary (CHO) cell lines were transfected with human *ASBT* and GAL4-NR and treated with DMSO (vehicle, *n* = 12), tCDCA (*n* = 3), B6 smvB BA pool (*n* = 4), or *Tnf^ΔARE/^*^+^ smvB BA pool (*n* = 4) for 24 hr. (B) Volcano plots of pulldown analyses for the distribution of mass features differentially enriched by RORγ. Cell lysates from *Escherichia coli* expressing human RORγ ligand-binding domain (LBD) or DMSO (vehicle) were incubated with B6 or *Tnf^ΔARE/^*^+^ smvB BA pools for 1 hr. Dashed lines represent significance cut-offs (*P*-value < 0.05, log_2_FC > 2). Masses of enriched BA species were highlighted. (C) Chemical structure and concentration of taurine-conjugated muricholic acids in smvB from B6 and *Tnf^ΔARE/^*^+^ mice. Chemical structure of tαMCA (*left*), tβMCA (*middle*), and tωMCA (*right*). (D) Flow cytometric analyses of IL17-A-GFP^+^ cells. CD4^+^CD25^-^CD44^-^ T cells were purified from IL-17A-GFP reporter mice, polarized, and treated with B6 smvB BA pool containing *Tnf^ΔARE/^*^+^ level of tβMCA (14.2 μM) or *Tnf^ΔARE/^*^+^ smvB BA pool containing B6 level tβMCA (103.6 μM) for 3 days (*n* = 6). IL-17A-GFP^+^ cells were gated on live CD4^+^ T cells. Representative flow plots (*left*), normalized frequency of IL-17A-GFP^+^ cells (*right*). (E and F) TR-FRET analyses of hRORγ LBD activation by BA pools. hRORγ LBD and FITC-labeled MED1/TRAP220 coactivator ID2 peptide were (E) titrated (*n* = 2) or (F) incubated (*n* = 12) with B6 smvB BA pool containing *Tnf^ΔARE/^*^+^ level of tβMCA (14.2 μM) or *Tnf^ΔARE/^*^+^ smvB BA pool containing B6 level of tβMCA (103.6 μM). BA pools (E) were titrated at the molar equivalent concentrations of 10^-3^×, 10^-2^×, 10^-1^× and 1×. (G) TR-FRET analyses of hRORγ LBD activation by individual BAs. hRORγ LBD and FITC-labeled MED1/TRAP220 coactivator ID2 peptide were titrated with indicated individual BAs (*n* = 2). (H) Flow cytometric analyses of IL-17A-GFP cells. CD4^+^CD25^-^CD44^-^ T cells were purified from IL-17A-GFP reporter mice, polarized, and treated with B6 smvB BA pool or B6 smvB pool lacking tβMCA (103.6 μM), tDCA (3.6 μM), or tLCA (0.19 μM) for 3 days (*n* = 8). IL-17A-GFP^+^ cells were gated on live CD4^+^ T cells. Representative flow plots (*left*), normalized frequency of IL-17A-GFP^+^ cells (*right*). (I) TR-FRET analyses of hRORγ LBD activation by tDCA or tLCA. hRORγ LBD and FITC-labeled MED1/TRAP220 coactivator ID2 peptide were incubated with the indicated bile acids (360 µM tβMCA, 30 µM tDCA, 1 µM tLCA) or DMSO with or without hRORγ LBD as vehicle and low signal controls, respectively (*n* = 13). (J) Representative bile acid binding modes from flexible side chain molecular docking of hRORγ LBD bound to bile acid agonists and antagonists compared to the crystallized binding mode of RORγ antagonist digoxin (PDB 3B0W). (K) Leu324 side chain shifts in the digoxin-bound RORγ LBD crystal structure (PDB 3B0W) and conserved among all bile acid antagonists. Data are presented as the mean ± SEM. Statistical significance was determined by Brown-Forsythe and Welch ANOVA with Dunnett’s T3 multiple comparisons test (A), one-way ANOVA with Tukey’s (D and G) post hoc tests, Brown-Forsythe and Welch (I) post hoc tests or Welch’s t-test (F). \**P* < 0.05, \*\**P* < 0.01, \*\*\**P* < 0.001, \*\*\*\**P* < 0.0001. See also **Figure S8** and **S9**.

To interrogate direct interactions between enterohepatically cycling BAs and RORγt, we incubated reconstituted wild type or *Tnf^ΔARE/^*^+^ enterohepatic BA pools with control or recombinant hRORγ LBD-expressing bacterial extracts; species capable of binding hRORγ LBD were resolved by mass spectrometry following affinity purification.^28^ The results revealed direct and differential binding of hRORγ LBD by BAs within wild type and *Tnf^ΔARE/^*^+^ enterohepatic pools; one or more taurine-conjugated muricholic acid (tMCA) species, as well as two taurine-conjugated secondary BAs (tDCA, tLCA), bound hRORγ LBD within the wild type enterohepatic BA pool, whereas only tDCA and tLCA retained hRORγ LBD-binding within the *Tnf^ΔARE/^*^+^ pool (**Figure 4B**). Loss of tMCA:RORγ-binding within the *Tnf^ΔARE/^*^+^ enterohepatic pool was consistent with marked depletion of muricholates (**Figure 4C**, **Figure S1A**) and suggested inhibitory interactions that may limit Th17 cell function in the healthy ileum. Conversely, tDCA and tLCA were quantitatively similar in both wild type and *Tnf^ΔARE/^*^+^ enterohepatic pools—relative abundance of these BAs increased in the *Tnf^ΔARE/^*^+^ pool given depletion of other species (**Figure 1B**, **Figure S1A**)—suggesting that tDCA and tLCA may augment Th17 activity during ileitis.

tMCAs include three stereoisomers that cannot be resolved by mass spectrometry absent internal standards—two 1° species (tαMCA, tβMCA) synthesized and conjugated in mouse hepatocytes, and one 2° species (tωMCA) produced via microbial metabolism in the LI and re-conjugation in the liver after intestinal absorption and portal recirculation.^29^ Of these, tβMCA has been shown to inhibit FXR with an IC_50_ of ∼40 μM.^3^ Assuming any putative tMCA:RORγ interaction carries a similar affinity, our data suggested that only tβMCA oscillates above and below this putative bioactive threshold in enterohepatic circulation of wild type and *Tnf^ΔARE/^*^+^mice, respectively (**Figure 4C**). To empirically test the contributions of each tMCA species to Th17 cell regulation by enterohepatic BA pools, we prepared otherwise identical wild type or *Tnf^ΔARE/^*^+^ enterohepatic (smvB) BA pools with or without wild type levels of tβMCA, or of all three tMCAs. The results confirmed that tβMCA was the main inhibitor of IL-17A^+^ Th17 cell development within mouse enterohepatic BA pools, as: (*i*) repression of IL-17A^+^ Th17 cell development by the wild type enterohepatic BA pool was partly alleviated by removal of tβMCA; (*ii*) normally unperturbed IL-17A^+^ Th17 cell development in the presence of *Tnf^ΔARE/^*^+^ BA pools decreased upon supplementation of this pool with wild type tβMCA levels; and (*iii*) further addition or removal of tαMCA/tωMCA had no additional impacts on IL-17A^+^ Th17 cell development (**Figure 4D, Figure S8M**). Notably, tβMCA alone did not suppress *in vitro* development of IL-17A^+^ Th17 cells (**Figure S8N-O**), suggesting that tβMCA may act primarily to limit RORγt activation by other co-circulating BA species.

To enable biochemical study of BA-RORγ interactions, we established time-resolved fluorescence resonance energy transfer (TR-FRET) assays in which cholesterol—a low-affinity endogenous RORγ agonist^24^—increased binding of purified hRORγ LBD protein to two MED1 coactivator peptides and decreased binding to a SMRT/*NCOR2* corepressor peptide (**Figure S9A**). Notably, *Med1* and Smrt/*Ncor2* were among the coregulators found by scRNA-seq to be expressed in ileal Th17 cells (**Figure S9B**). We next used these assays to examine tβMCA contributions to direct hRORγ LBD regulation by wild type and *Tnf^ΔARE/^*^+^ enterohepatic BA pools. Consistent with cell culture results, removal of tβMCA from the wild type enterohepatic BA pool increased hRORγ LBD binding to MED1 coactivator peptides, whereas addition of wild type tβMCA concentrations to the *Tnf^ΔARE/^*^+^ enterohepatic pool reduced this interaction (**Figure 4E**-**F**). At the same time, tβMCA alone did suppress basal hRORγ LBD binding to MED1 and high concentrations of tβMCA displaced SMRT corepressor peptides from hRORγ LBD (**Figure 4G**, **Figure S9C**), suggesting tβMCA acts as an RORγt antagonist, not an inverse agonist.

In contrast to tβMCA, cell culture and TR-FRET studies revealed that both tDCA and tLCA operate within enterohepatic BA pools as direct RORγt agonists. First, tDCA and tLCA each acted alone and in dose-dependent manners to increase *in vitro* IL-17A^+^ Th17 cell development and hRORγ LBD binding to MED1 coactivator peptides (**Figure 4G, Figure S8N**-**O**). Further, increased development of IL-17A^+^ Th17 cells in the presence of tβMCA-deficient wild type enterohepatic BA pools required presence of tDCA or tLCA (**Figure 4H**). Most importantly, tβMCA concentrations present in enterohepatic circulation of wild type, but not *Tnf^ΔARE/^*^+^, mice inhibited tDCA- or tLCA-mediated hRORγ LBD recruitment of MED1 coactivator peptides (**Figure 4I**, **Figure S9D**). As a whole, these findings suggest that co-circulating agonist (tDCA, tLCA) and antagonist (tβMCA) BA ligands are in direct competition for RORγt-binding in the ileal mucosa.

To gain insight into the structural logic behind BA-mediated RORγ agonism *vs*. antagonism, we used flexible side chain molecular docking. tDCA, tLCA and tβMCA each adopted an RORγ-binding pose similar to digoxin, a known antagonist previously crystallized with hRORγ LBD that contains a BA-like steroidal cholane scaffold (**Figure 4J**).^30,31^ For each BA, the taurine moiety adopted similar positioning within the hRORγ orthosteric ligand binding pocket as the furanone group of digoxin (**Figure 4J**). However, whereas BA agonists (tDCA, tLCA) pulled the Leu324 side chain on helix 3 away from the helix 12/AF-2 surface, antagonists (digoxin, tβMCA) shifted the Leu324 side chain towards the helix 12/AF-2 coregulator interaction surface (**Figure 4K**). In these models, repositioning of the Leu324 side chain towards side chains of Tyr502 and Phe506 on helix 12 involved C-6/C-7 hydroxyl groups in tβMCA and promoted unfavorable hydrophobic-hydrophobic interactions between H479, F486, F506 and Y502 in the RORγ LBD that are necessary for transcriptional activation.^32^ Conversely, hydrophobic methylene groups in tDCA and tLCA interacted favorably with the Leu324 side chain, drawing it towards the BA steroid ring and away from the helix 12/AF-2 surface, which left this hydrophobic network intact.

Additional studies revealed that BA conjugation with taurine—and to a lesser extent glycine—increased the potency with which DCA and LCA bound and activated hRORγ LBD (**Figure S9E-H**). These results were important, as they provided a possible molecular explanation for why taurine-conjugated BA species outcompeted (in some cases more abundant) unconjugated counterparts for hRORγ LBD-binding within complex enterohepatic BA pools (**Figure 4B**). Flexible side chain molecular docking studies suggested that the sulfonic acid group of tDCA and carboxylic acid groups in DCA and gDCA form hydrogen bonds/polar interactions with backbone amide groups of Gln286 and/or Leu287. However, progressively lengthened C-24 side chains in gDCA and tDCA: *(i*) contain an acetamide group that enables polar interactions with the Gln286 backbone amide and carboxamide side chain in RORγ; (*ii*) increasingly position the cholane steroid nucleus closer to the helix 12/AF-2 surface; and (*iii*) form additional polar interactions with residues on hRORγ helix 3 (Cys320, Leu324, His323) (**Figure S9I**). At the same time, numerous other taurine-conjugated 1° and 2° BA species present in human and mouse enterohepatic pools (tCA, tCDCA, tUDCA) did not interact with hRORγ LBD (**Figure 4G**), suggesting that both modifications to the BA steroid ring and amino acid conjugation contribute to specifying BA-RORγ interactions.

### tβMCA supplementation restores regulation of ileal Th17 cells in *Tnf^ΔARE/+^* mice

As tβMCA is an RORγt antagonist whose depletion from enterohepatic circulation in *Tnf^ΔARE/^*^+^ mice is associated with Th17-associated ileitis, we reasoned that replenishing tβMCA enterohepatic levels in *Tnf^ΔARE/^*^+^ mice might re-assert BA regulation over ileal Th17 cells. To begin testing this, we first determined oral dosing requirements needed to restore wild type levels of enterohepatically circulating tβMCA in *Tnf^ΔARE/^*^+^ mice (**Figure 5A**). Younger (13-week-old) *Tnf^ΔARE/^*^+^ mice were used for these studies, as: (*i*) tβMCA requires active (Asbt-mediated) transport for efficient ileal absorption;^33^ (*ii*) chronic ileitis reduces Asbt/*Slc10a2* expression (**Figure 2**); and (*iii*) increases in ileal Th17 cells and neutrophils were already apparent in *Tnf^ΔARE/^*^+^ mice around this timepoint (**Figure 3D, 3H**). Up to 10 mg of oral tβMCA was tolerated, in line with prior reports,^3,34^ but 1 mg was sufficient to increase tβMCA levels in enterohepatic circulation (smvB) of *Tnf^ΔARE/^*^+^ mice to those measured in age-matched wild type control animals (**Figure 5B**). A lower dose (0.2 mg) had no effect on enterohepatic tβMCA levels in *Tnf^ΔARE/^*^+^ mice, compared with untreated *Tnf^ΔARE/^*^+^ counterparts, and the 10 mg dose increased tβMCA to supraphysiological levels in both enterohepatic circulation (smvB) and the LI (cecum) (**Figure 5B**-**C**). These results confirm that orally delivered tβMCA is efficiently absorbed in the ileum and incorporated into enterohepatic circulation, even in *Tnf^ΔARE/^*^+^ mice where ileitis reduces and remodels enterocyte-mediated ileal BA uptake.

**Figure 5.**
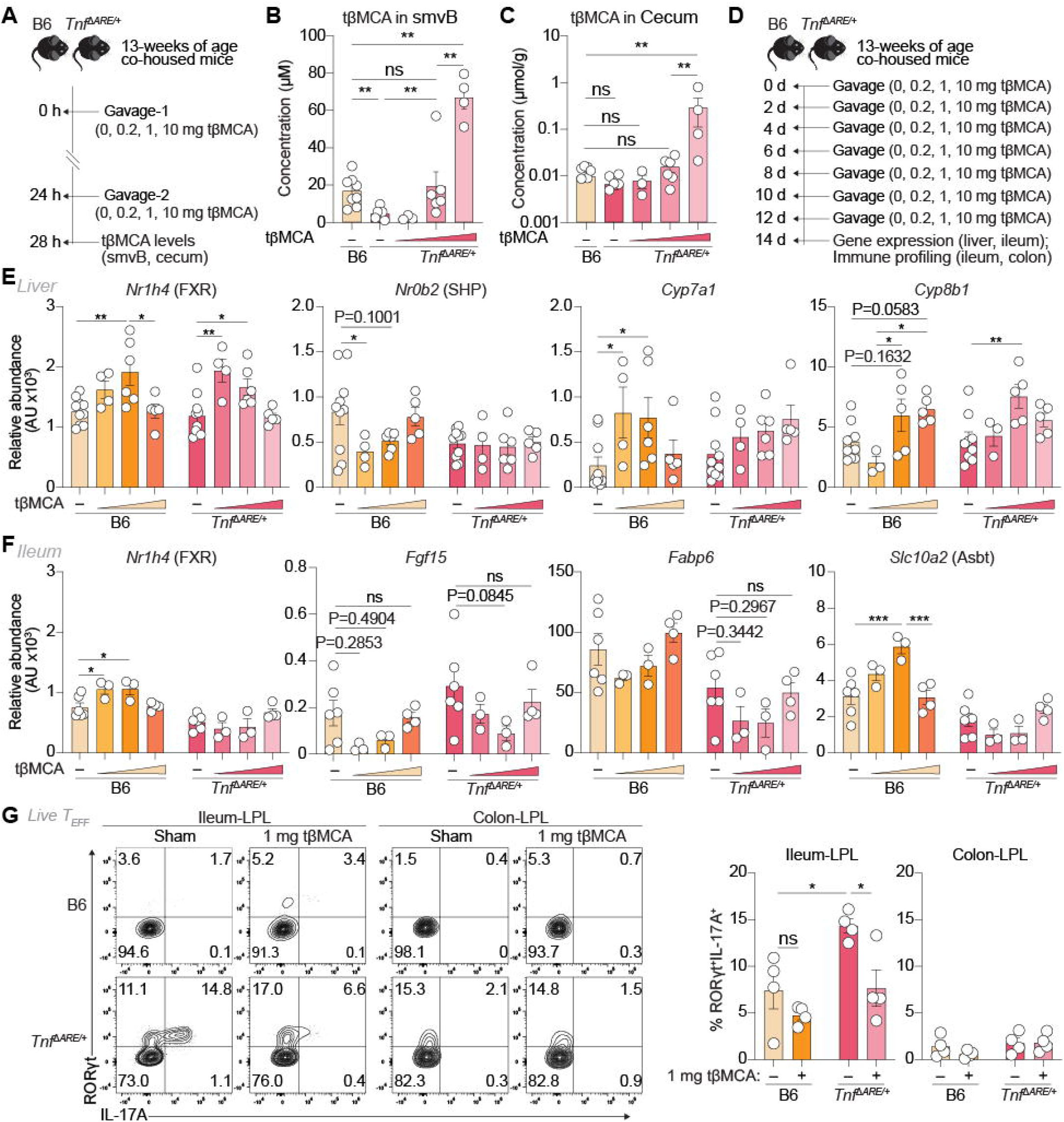
Restoring healthy enterohepatic levels of tβMCA represses ileal Th17 cells in *Tnf*^ΔARE/+^ mice. (A) Schematic of acute tβMCA treatment. 13-week-old B6 and *Tnf^ΔARE/^*^+^ mice were co-housed at weaning and treated with 0, 0.2, 1, or 10 mg/dose of tβMCA by oral gavage at 0 and 24 hr, smvB was collected after fasting for 4 hr. (B and C) Concentration of tβMCA in smvB (B) and cecum (C) following acute tβMCA treatment (*n* = 3-8 mice). (D) Schematic of two-week tβMCA treatment. 13-week-old B6 and *Tnf^ΔARE/^*^+^ mice were co-housed at weaning and treated with 1 mg/dose of tβMCA by oral gavage every other day for two weeks (*n* = 3-10 mice). (E and F) mRNA expression of BA-metabolizing genes in the liver (E) and ileum (F) after two-week tβMCA treatments. (G) Frequencies of IL-17A-producing Th17 cells in ileal and colonic lamina propria. Representative flow plots (*left*), frequency of RORγt^+^IL-17A^+^ (*right*). RORγt^+^IL-17A^+^ cells were gated on Live CD45^+^TCRβ^+^CD4^+^CD44^hi^CD62L^-^Foxp3^-^ cells. 13-week-old B6 and *Tnf^ΔARE/^*^+^ mice were co-housed at weaning and treated with PBS (Sham) or 1 mg/dose of tβMCA by oral gavage every other day for two weeks (*n* = 4-6 mice). Data are presented as the mean ± SEM. Statistical significance was determined by one-way ANOVA with Tukey’s post hoc test (B, C) and two-way ANOVA with Tukey’s post hoc test (E to G). \**P* < 0.05, \*\**P* < 0.01, \*\*\**P* < 0.001. See also **Figure S10**.

Having established dosing requirements, we next performed longer-term treatment studies to examine *in vivo* Th17 cell regulation by orally delivered tβMCA in *Tnf^ΔARE/^*^+^ mice. For this, we administered 1 mg oral tβMCA to co-housed wild type or *Tnf^ΔARE/^*^+^ mice every other day for 2 weeks (**Figure 5D**); a known pharmacodynamic effect of tβMCA (inhibition of FXR^3^) was measured and related to regulation of RORγt-mediated Th17 cell function in the ileum and colon. The 2-week treatment period was used to enrich for more proximal effects of tβMCA administration and avoid accumulation of less direct downstream phenotypes. As expected, mice receiving oral tβMCA displayed changes in liver and ileum gene expression consistent with FXR antagonism, including: (*i*) reduced hepatic expression of the FXR target gene, SHP/*Nr0b2*; (*ii*) increased hepatic expression of rate-limiting BA biosynthetic enzymes repressed by SHP (*Cyp7a1*, *Cyp8b1*);^35,36^ (*iii*) lower ileal expression of *Fgf15* and *Fabp6*, two genes induced by FXR in enterocytes; and (*iv*) higher ileal expression of Asbt/*Slc10a2*, which as noted above is repressed by FXR activity in healthy enterocytes (**Figure 5E**-**F**). Several of these effects were not apparent in *Tnf^ΔARE/^*^+^ mice, consistent with baseline FXR dysregulation, although *Tnf^ΔARE/^*^+^ mice treated with tβMCA did show reduced ileal *Fgf15* and increased hepatic *Cyp8b1* expression (**Figure 5E**-**F**).

*Tnf^ΔARE/^*^+^ mice receiving 1 mg oral tβMCA also displayed reduced frequencies of IL-17A^+^ Th17 cells in the ileum, compared with untreated controls, and levels of ileal Th17 cells in tβMCA-treated *Tnf^ΔARE/^*^+^ mice approached baseline Th17 cell levels observed in untreated wild type controls (**Figure 5G**). Notably, effects of tβMCA on ileal Th17 cells were specific, as no other changes in numbers or percentages of many other immune cell lineages, including neutrophils, were observed in ileums of tβMCA-treated *Tnf^ΔARE/^*^+^ mice, and tβMCA treatment did not impact Th17 cell numbers or frequencies in the colon, again supporting a local mechanism of RORγt antagonism that follows active ileal uptake by enterocytes (**Figure 5G, Figure S10A-C**). Together, these data not only confirm that tβMCA can inhibit RORγt-mediated Th17 cell function *in vivo*; they establish that aberrant enterohepatic BA pools generated during ileitis can be corrected using rationally designed oral supplements to restore BA and immune homeostasis.

## Discussion

By mapping complex endogenous BA pools that circulate enterohepatically through the healthy or inflamed ileal mucosa of mice—and establishing new workflows that permit discovery of discrete BA-NR interactions within these pools—we report here that BA-dependent ileal immune regulation involves competitive binding of RORγt by two agonist (tDCA, tLCA) and one antagonist (tβMCA) BA species. These insights are important because they move beyond contemporary models of single BA species acting in isolation and towards a more physiological ground-truth, where functions of gut mucosal immune cells are shaped by the absolute and relative abundance of dozens-to-hundreds of pharmacodynamically unique BA species circulating together in composite pools. Competitive and antagonistic BA interactions have been described previously for FXR, which in ileal enterocytes and hepatocytes is activated by tCDCA but inhibited by tβMCA.^2,3^ Our results show that competition between BAs for RORγt also shapes immunoregulatory outcomes. Thus, it is reasonable to expect that intra-pool competitive dynamics are the rule, not the exception, by which BAs modulate host physiology *in vivo*.

Functional outcomes of competitive and mutually antagonistic BA interactions with single NRs should be determined by two main variables: concentrations and binding affinities (K_d_). In the case shown here for RORγ, fitting of TR-FRET curves suggest that the two RORγ agonist ligands (tDCA and tLCA) bind hRORγ LBD with EC_50_ values (which approximate K_d_) of ∼9 μM and ∼0.5 μM, respectively. Conversely, tβMCA inhibits hRORγ LBD-mediated coactivator recruitment with a K_i_ of ∼130 μM. In isolation, 10- to 20-fold higher affinities of agonist (tDCA and tLCA) *vs*. antagonist (tβMCA) BA ligands for RORγt could suggest that exposure of ileal Th17 cells to enterohepatically cycling BA pools generally favors RORγt agonism. However, our result also show that the mean steady-state concentration of enterohepatically circulating tβMCA in healthy 30-week-old wild type mice (103.6 μM) approaches ∼80% of its K_i_ value for RORγ LBD. By contrast, steady-state concentrations of tDCA (3.6 μM) or tLCA (0.19 μM) represent only ∼40% of their respective K_d_ values for RORγ LBD. Ileitis in *Tnf^ΔARE/^*^+^ mice depletes enterohepatic tβMCA levels nearly 10-fold (to 14.2 μM) but leaves tDCA and tLCA more intact (2.4 μM and 0.14 μM, respectively). Thus, while other variables (*e.g*., BA retention in ileal tissue, feeding *vs*. fasting, etc.) will also impact precise intracellular concentrations of these and other BAs within ileal immune cells *per se*, these results support a model in which the ileal BA environment in healthy mice is generally restrictive to RORγt-mediated Th17 cell function, in contrast to one that is more favorable to Th17 cells in *Tnf^ΔARE/^*^+^ mice with ileitis. These results also raise the possibility that a pathogenic feed-forward loop exists between aberrant BA enterohepatic circulation and inflammatory T cell function during ileitis, where inflammatory Th17 cells are functionally amplified by the same aberrant BA pools they help evoke.

Our work sheds new light on the complex roles of BAs in CD, which have been incompletely understood and controversial. On one hand, ileal CD has long been associated with BA malabsorption (BAM), ^37^ which has prompted speculation that BAs must be anti-inflammatory in CD (*i.e*., because their ileal absorption is reduced in patients).^38^ In a general sense, our results confirm BAM is a conserved feature of ileitis in both human CD patients and *Tnf^ΔARE/^*^+^ mice. However, our data also establish that the enterohepatic BA pool generated during ileitis is not only smaller; it is also compositionally different and enriched for pro-inflammatory features. In particular, ileitis-associated BA pools are disproportionately depleted of liver-derived 1° cBA (which in mice include the RORγt antagonist, tβMCA) but enriched in microbially generated 2° uBA and 2° metabolites. Our finding that compositionally distinct enterohepatic BA pools have divergent immunoregulatory functions adds an important new layer to how BAM is contextualized in CD and draws parallels with IBD-associated microbial dysbiosis, which is not simply expressed in terms of total biomass (*i.e*., more or fewer bacteria), but rather by changes in the composition and metabolic function of pro- and anti-inflammatory community members.

Our study adds three novel BA-NR interactions to a still-growing list. In the colon, alloiso- LCA has been shown to stabilize Foxp3 expression by activating NR4A1 during peripheral (p)Treg cell development, whereas LCA activation of VDR promotes IL-10 production in mature pTreg cells.^39,40^ In parallel, 3-oxo-LCA and iso-LCA inhibit colonic Th17 cell function via direct RORγt antagonism.^25^ It is notable that our studies of the enterohepatic BA pool identified tDCA, tLCA and tβMCA—but not 3-oxo-LCA or iso-LCA—as physiological RORγt ligands in the ileum. Most likely, this is because of extreme compositional differences between BA pools in the small *vs*. large intestines. Indeed, our results show that the pool of smvB BAs which circulates enterohepatically through the ileal mucosa of wild type B6 mice is ∼7x larger than the pool that transits the colonic mucosa and enters imvB (∼310 μM vs. ∼44 μM). Further, whereas ∼60% of the ileal BA pool in wild type mice is comprised of liver-derived 1° cBAs, >60% of colonic BAs are microbially produced 2° uBA species and metabolites.^4^ Given these marked differences in the size and complexity of ileal and colonic BA pools, differences in BA-dependent immune regulation between these two intestinal segments stand to reason. Future efforts to empirically define shared *vs*. unique mechanisms of BA-mediated immune regulation in the ileum and colon will have important implications in the understanding and treatment of IBDs that involve the large *vs*. small bowel.

Discovery of tβMCA as a physiological RORγ antagonist in mice has important therapeutic implications for human CD. Although tβMCA is only present endogenously in mice (not humans), there is precedent for BAs found in other animals leading to human therapeutics. Ursodeoxycholic acid (UDCA), for example, is a minor 2° BA in humans that was first identified as an abundant 1° BA in bears; UDCA (*a.k.a*., Ursodiol) is now standard-of-care therapy for cholestatic liver diseases, including primary biliary cholangitis and intrahepatic cholestasis of pregnancy and is used as adjunct therapy for biliary atresia.^41^ Thus, it may be possible to explore clinical utility of tβMCA in human CD. At the same time, the fact that tβMCA inhibits both RORγ and FXR is likely to limit its therapeutic utility in CD; FXR function is generally reduced in IBD patients and FXR agonists are being explored as IBD therapeutics.^42–44^ Thus, tβMCA may prove most useful as a chemical scaffold to discover new semi-synthetic small molecules that preserve efficient ASBT-mediated ileal targeting, but that show enhanced potency and selectivity for RORγ over FXR. Whether tβMCA itself, a semi-synthetic tβMCA derivative or another BA altogether, our study suggests that rational approaches to quantify and correct ileitis-associated compositional shifts within the enterohepatic BA pool could be a viable approach to restore BA and immune homeostasis in human CD. This is especially important given the current lack of therapeutic options for patients with mild-to-moderate CD, and the hesitancy of these patients to initiate biologic therapy early.^45^ Rationally designed BA-based oral supplements could help fill this gap and provide new tools for disease ‘interception’ in CD.^46^

### Limitations

This study describes mechanisms of BA-mediated immune regulation in mice. Parallel analysis of serum BA signatures in human CD patients *vs*. controls suggests ileitis is likely to produce generally similar perturbations to the pools of enterohepatically cycling BAs in humans (*i.e*., reduced levels of liver-derived 1° cBAs, proportional enrichment of microbially generated 2° uBA species and metabolites). However, species-specific differences in hepatic and gut microbial BA metabolic pathways establish that the precise composition of ileitis-associated BA pools in human CD patients and *Tnf^ΔARE/^*^+^ mice will be different. In addition, human livers conjugate 1° BAs with mostly glycine, whereas taurine is used in mice. As such, it will be important for future studies to compare and contrast immunoregulatory functions of mouse and human enterohepatic BA pools, although defining human enterohepatic BA pools for functional studies will not be straightforward. For example, the pool of smvB BAs which circulates enterohepatically through the ileal mucosa is both quantitatively and functionally distinct from other pools measured in more accessible tissues of the same animals (*e.g*., serum, stool; **Figure S8A**). Thus, functional studies of *bona fide* enterohepatic BA pools in humans will either require direct measurement of smvB BAs, or development of new computational approaches to predict quantitative features of human enterohepatic BA pools from serum BA profiles.

## Acknowledgements

The authors thank Core Facility staff at Dartmouth-Hitchcock Medical Center (DHMC), Dartmouth Cancer Center and Herbert Wertheim UF Scripps Institute for Biomedical Innovation and Technology for technical support. We also acknowledge support from the Analytical Chemistry Core at Harvard Medical School (https://analyticalchem.hms.harvard.edu/) for targeted analysis of tβMCA following oral supplementation. **Funding:** This research was supported by startup funds from the Walter and Carole Young Center for Digestive Health at Dartmouth Health (M.S.S) or Vanderbilt University (D.J.K.), sub-awards from the Dartmouth Cystic Fibrosis Research Center grant P30DK117469 and the Dartmouth Cancer Center grant P30CA023108 (M.S.S), a Cystic Fibrosis Foundation grant OTOOLE22G0 (G.A.O.), internal funds from the University of Miami through a philanthropic gift, and by the Leona M. and Harry B. Helmsley Charitable Trust grant 2004-03822 (M.T.A) and grants from Abbvie, Celltrion, Johnson & Johnson, Lilly, Pfizer and Takeda (C.A.S). Additional funding was provided by National Institute of Health (NIH) grants R01AI143821, R01AI164772, U01AI163063, and R01DK145055 (to M.S.S), R01ES033988 (to G.A.O.), and R01DK140485 (to P.A.D). scRNA-seq was supported by Dartmouth Center for Quantitative Biology (CQB), funded in part by NIH grants P20GM130454 and S10OD025235. The Center for Clinical Genomics and Advanced Technology in the Department of Pathology and Laboratory Medicine of the Dartmouth Hitchcock Health System, which includes the Pathology Shared Resource, at the Dartmouth Cancer Center is supported by NCI Cancer Center grants 5P30CA023108-37 and RRID SCR-023479. DartLab is supported by the NCI Cancer Center Support grant P30CA023108 and RRID SCR-019165.

## Author contributions

Study design: Z.X., K.S., D.S., P.A.D., D.J.K., and M.S.S. Data generation: Z.X., K.S., A.D-E., J.L., P.M-T., F.N., A.D.H., H.Y, B.J.H., and A.R-P. Bioinformatics: Z.X., H.D.D., A.B., S.S. Manuscript: Z.X., K.S., J.L., C.L.H., E.M.T., I.K.K., C.A.S., G.A.O., C.T.W., H.M.K., M.T.A., P.A.D., D.J.K., and M.S.S. Principal Investigators: C.A.S., G.A.O., D.S., F.C., H.M.K., M.T.A., P.A.D., D.J.K., and M.S.S.

## Supplemental information

Supplemental information for this manuscript includes supplemental figures (Figure S1-S10) and one supplemental table (Table S1).

---

**Figure S1. Ileitis reshapes the landscape of bile acid pools, related to Figure 1**

(A) Triangle plot of BA concentrations in siLC, smvB, feces, and pB from co-housed B6 and *Tnf^ΔARE/^*^+^ mice. Data presented are the median concentration of BA species from B6 mice (*bottom left*) and *Tnf^ΔARE/^*^+^ mice (*top right*).

(B) Schematic of enterohepatic BAs absorption in the ileum.

(C) β-diversity of cecal microbial analyses in B6 and *Tnf^ΔARE/^*^+^ mice.

(D) Relative abundance of BA-metabolizing bacteria in B6 and *Tnf^ΔARE/^*^+^ mice. All mice were analyzed at 30-weeks of age (*n* = 7 mice).

Data are presented as the mean ± SEM. Statistical significance was determined by Wilcoxon signed-rank test. ns, not significant.

**Figure S2. Serum bile acid profiles in healthy controls and Crohn’s disease patients, related to Figure 1**

(A) Concentrations of BA species in serum from healthy controls (HC) and Crohn’s diseases patients. Data presented are the median concentration of BA species from HC (*bottom left*) and ileal Crohn’s diseases patients (iCD, *top right*).

(B) Total BA concentrations in serum from HC and iCD.

(C) PCA analysis of BA species in serum from HC and iCD. The top five principal components of BA species are shown. 17 pairs of HC and iCD housemates were analyzed.

Data are presented as the mean ± SEM. Statistical significance was determined by two-tailed paired Wilcoxon test (B) and Wilcoxon signed-rank test (C). \**P* < 0.05, \*\**P* < 0.01.

**Figure S3. Characterization of *Slc10a2*-expressing cells in *Tnf^ΔARE/^*^+^ileitis, related to Figure 2**

(A) Protein expression of *Slc10a2* in terminal ileum from B6, *Slc10a2*-deficient, and *Tnf^ΔARE/^*^+^ mice. Representative images of immunohistochemistry staining (*left*), scale bar, 100 μm. Quantification of % area of *Slc10a2* expression (*right*). Data were quantified from two or three independent sections per mouse (*n* = 2-3) with 5-6 fields per section.

(B) Schematic of scRNA-Seq of ileal epithelial fractions. B6, *Slc10a2*^-/-^, *Tnf^ΔARE/^*^+^, and *Tnf^ΔARE/^*^+^;*Slc10a2*^-/-^ were co-housed at weaning age until 15-weeks of age. 2 mice poured per sample. Ileal epithelial fractions were obtained and purified after dead cells removal by MACS purification.

(C) Dot plots of feature gene expression in each cluster of ileal epithelial fractions.

(D) UMAP of absorptive enterocytes subclusters and BA-metabolizing genes expression.

(E) Heatmap of top 20 differentially expressed genes in absorptive enterocytes subclusters. *P*_adj_ < 0.05 with descending log_2_Fold Change (log_2_FC).

(F) Volcano plot of differentially expressed genes between Cluster 1 and Cluster 0 in absorptive enterocytes subclusters. *P*_adj_ < 0.05, log_2_ (FC) > 0.5.

(G) Volcano plot of differentially expressed genes between Cluster 1 and Cluster 3 in absorptive enterocytes subclusters. *P*_adj_ < 0.05, log_2_ (FC) > 0.5.

Statistical significance was determined by one-way ANOVA with Tukey’s post hoc test (A) and Wilcoxon signed-rank test (D and E). \**P* < 0.05, \*\**P* < 0.01, \*\*\**P* < 0.001, \*\*\*\**P* < 0.0001.

**Figure S4. scRNA-seq of *Slc10a2*-expressing enterocytes in human Crohn’s disease, related to Figure 2**

(A) Dot plot of feature gene expression of cell clusters in human scRNA-Seq dataset. EECs, Endothelium cells.

(B) Dot plot of feature gene expression in epithelial cells subclusters. SCs, stem cells; TAs, transit-amplifying epithelial cells; Entero, Enterocytes; GCs, Goblet cells.

(C) UMAP of ileal epithelial cells subclusters and BA-metabolizing genes.

(D) Volcano plot of differentially expressed genes in Cluster 6. *vs.* Cluster 0 (*left*) and Cluster 6. *vs.* Cluster 14 (*right*) in ileal epithelial sub-clustering. *P*_adj_ < 0.05, log_2_ (FC) > 0.25.

Statistical significance was determined by Wilcoxon signed-rank test.

**Figure S5. BA pools ablation alleviates ileal inflammation in *Tnf^ΔARE/^*^+^ileitis, related to Figure 3**

(A) BA concentrations in feces (*n* = 4 mice).

(B) Representative H&E staining of terminal ileum (*n* = 3-9 mice). Representative images (*left*, Scale bar, 100 μm), quantification of inflammatory scores (*right*). 25-week-old B6 and *Tnf^ΔARE/^*^+^ mice were co-housed at weaning and fed with 2% dietary cholestyramine for 5 weeks (A and B).

(C) Representative H&E staining of terminal ileum (*n* = 3-13 mice). Representative images (*left*, Scale bar, 500 μm), quantification of inflammatory scores (*right*).

(D) Microbial analyses between *Tnf^ΔARE/^*^+^. *vs. Tnf^ΔARE/^*^+^;*Slc10a2*^-/-^ mice (*n* = 7 mice). α-diversity (*left*), β-diversity (*right*). B6, *Slc10a2*^-/-^, *Tnf^ΔARE/^*^+^, and *Tnf^ΔARE/^*^+^;*Slc10a2*^-/-^ mice were co-housed at weaning and analyzed at 30-weeks of age.

(E) Relative mRNA expression of SFB in feces. SFB, Segmented filamentous bacteria (*n* = 7 mice). B6, *Slc10a2*^-/-^, *Tnf^ΔARE/^*^+^, and *Tnf^ΔARE/^*^+^;*Slc10a2*^-/-^ mice were co-housed at weaning and analyzed at 15-weeks of age. Positive control (Pos) was obtained from mice colonized with SFB.

Data are presented as the mean ± SEM. Statistical significance was determined by one-way ANOVA with Šídák’s post hoc test (A and B), one-way ANOVA with Tukey’s post hoc test (C and E) and Wilcoxon signed-sum test (D). \**P* < 0.05, \*\**P* < 0.01, \*\*\**P* < 0.001, \*\*\*\**P* < 0.0001, ns, not significant.

**Figure S6. scRNA-Seq profiling of CD4^+^ T cells from ileal lamina propria in *Tnf^ΔARE/^*^+^ileitis, related to Figure 3**

(A) Schematic of scRNA-Seq of immune cells from ileal lamina propria. B6, *Slc10a2*^-/-^, *Tnf^ΔARE/^*^+^, and *Tnf^ΔARE/^*^+^;*Slc10a2*^-/-^ were co-housed at weaning until 15-weeks of age, CD4^+^ T cells (TCRβ^+^CD4^+^) and non-CD4^+^ T cells (CD4^-^) were sorted and mixed (1:1) for scRNA-Seq. Two mice were pooled per sample.

(B) Violin plot of *Cd3e* expression across all identified clusters.

(C) Dot plot of marker gene expression across cell clusters.

**Figure S7. BA pools modulate local immune function in *Tnf^ΔARE/^*^+^ileitis, related to Figure 3**

(A) Gating strategy for T cells and innate lymphocytes.

(B) Percentages of T cells and innate lymphocytes in ileum and colon. RORγt^+^Tregs, Th17 cells, Th1 cells, GATA3^+^Tregs and Th2 cells were gated on lineage (EpCAM, CD11b, CD19, F4/80, Ly6G and Ly6C)^-^CD45^+^TCRβ^+^CD4^+^ CD44^hi^ cells. ILC2 and ILC3 were gated on lineage^-^CD45^+^TCRβ^-^TCRγδ^-^CD3e^-^NK1.1^-^CD90.2^+^CD127^+^ cells. RORγt^+^TCRγδ^+^ cells were gated on lineage^-^CD45^+^TCRγδ^+^ cells.

(C) Gating strategy for T cells, B cells and myeloid cells.

(D) *t*-SNE plots and percentages of immune populations in ileum and colon. B6, *Slc10a2*^-/-^, *Tnf^ΔARE/^*^+^, and *Tnf^ΔARE/^*^+^;*Slc10a2*^-/-^ were co-housed at weaning and analyzed at 15-weeks of age (*n* = 9 mice; B and D) or 30-weeks of age (*n* = 3 mice; B). A total of 7,000 CD45^+^ cells per group (ileum) and 10,000 CD45^+^ cells per group (colon) were analyzed, all populations were calculated as a proportion of CD45^+^ cells.

Data are presented as the mean ± SEM. Statistical significance was determined by one-way ANOVA with Šídák’s post hoc test. \**P* < 0.05, \*\**P* < 0.01, ns, not significant.

**Figure S8. BA pools regulate Th17 cells differentiation *in vitro*, related to Figure 4**

(A) Flow cytometric analyses of IL-17A-GFP^+^ cells from cells treated with smvB, pB and imvB BA pools of B6 mice. Cells were treated with BA pools in smvB, pB and imvB of B6 mice for 3 days (*n* = 4), BA pools were reconstituted as indicated in Table. S1.

(B) Cell viability of total CD4^+^ T cells. Cells were treated with smvB BA pools of B6 and *Tnf^ΔARE/^*^+^ mice for 3 days (*n* = 10).

(C and D) Proliferation of Th17 cells. Cells were treated with smvB BA pools of B6 and *Tnf^ΔARE/^*^+^ mice for 3 days (*n* = 4).

(E to I) mRNA expression of lineage-associated transcription factors and cytokines in day 3-polarized non-pathogenic Th17 cells (npTh17; E), pathogenic Th17 cells (pTh17; F), Th1 cells (G), Th2 cells (H), or iTreg cells (I). Cells were activated and expanded in the presences of polarizing cytokines and reconstituted smvB BA pools (*n* = 6).

(J) Kinetics of phosphorylation of STAT3 (pSTAT3). Cells were treated with smvB BA pools of B6 and *Tnf^ΔARE/^*^+^, MFI of pSTAT3 were analyzed at 0, 0.5 hr, 2 hr, 6 hr, 12 hr, and 24 hr (*n* = 3).

(K and L) Kinetics of RORγt expression (F) and IL-17A-GFP^+^ cells (G). Cells were treated with smvB BA pools of B6 and *Tnf^ΔARE/^*^+^, MFI of RORγt and percentage of IL-17A-GFP^+^ cells were analyzed at 6 hr, 24 hr, 48 hr, 66 hr and 72 hr (*n* = 3).

(M) Flow cytometric analyses of IL-17A-GFP^+^ Th17 cells (*n* = 7) cultured for 3 days in the presence of reconstituted smvB BA pools mimicking wild type or *Tnf^ΔARE/^*^+^ mice containing or lacking B6 levels of taurine conjugated muricholic acids (tβMCA, tαMCA, tωMCA).

(N and O) Day 3 cell viability (M) or IL-17A-GFP^+^ cells (O) in cultured Th17 cells treated with titrating concentrations of tβMCA, tLCA or tDCA only.

Data are presented as the mean ± SEM. Statistical significance was determined by one-way ANOVA with Tukey’s post hoc test (A to D, H to L, M), two-way ANOVA with Tukey’s post hoc test (E to G). \**P* < 0.05, \*\**P* < 0.01, \*\*\**P* < 0.001, \*\*\*\**P* < 0.0001, ns, not significant.

**Figure S9. tβMCA directly competes against tLCA/tDCA in RORγ activity, related to Figure 4**

(A) Screening of coregulator against hRORγ LBD using the weak RORγ partial agonist cholesterol. 21 FITC-labeled coregulator peptides derived from nuclear receptor co-activators and co-repressors (*n* = 2) were used. EC_50_/IC_50_ values for select conditions are shown in parenthesis with fitted error ranges in square brackets.

(B) Dot plot of coregulators expression in Th17 cells. scRNA-Seq was performed as indicated in Fig. S6. (C) Dose-dependent effects on interaction between tβMCA and RORγ co-repressor ID2 peptide. TR-FRET assay was performed using hRORγ LBD and FITC-labeled SMRT/*Ncor2* co-repressor ID2 peptide titrated with tβMCA (*n* = 2).

(D) Competitive interaction of tβMCA and tLCA with RORγ co-activator ID2 peptide. TR-FRET assay was performed using hRORγ LBD and FITC-labeled MED1/TRAP220 coactivator ID2 peptide in competition mode. Antagonist mode: tβMCA, tαMCA and tωMCA were titrated in the presence of a half-saturating amount of 1 µM tLCA agonist.

(E and G) RORγ activities of individual ligands within the BA pools. LCA and t/gLCA (E); DCA and t/gDCA (G). TR-FRET assay was performed using hRORγ LBD and FITC-labeled MED1/TRAP220 co-activator ID2 peptide titrated with the indicated individual bile acids (*n* = 2).

(F and H) Thermostability curve of RORγ-binding for tLCA (F) and tDCA (H).

(I) Flexible side chain molecular docking models of hRORγ LBD bound to unconjugated and taurine/glycine-conjugated DCA and LCA. Representative bile acid binding modes show that glycine- and taurine-conjugated BAs forms additional polar interactions with residues lining the hRORγ LBD orthosteric ligand-binding pocket and extend longer modified steroid cholane scaffolds deeper in the orthosteric pocket towards the helix 12/AF-2 coregulator interaction surface.

Data are presented as the mean ± SEM.

**Figure S10. tβMCA regulates immune profile in *Tnf^ΔARE/^*^+^ileitis, related to Figure 5**

(A and B) *t*-SNE plots (A) and percentages (B) of immune populations in ileum and colon at 15-weeks of age. A total of 5,600 CD45^+^ cells per group (ileum) and 1,500 CD45^+^ cells per group (colon) were analyzed, all populations were calculated as a proportion of CD45^+^ cells.

(C) Percentages of T cells and innate lymphocytes in ileum and colon. 13-week-old B6 and *Tnf^ΔARE/^*^+^ mice were co-housed at weaning and treated with PBS or 1 mg/dose of tβMCA every other day for 2 weeks by oral gavage (*n* = 4-6 mice). Cell populations were analyzed via the gating strategy indicated in Figure S7A and S7C. Data are presented as the mean ± SEM. Statistical significance was determined by two-way ANOVA with Tukey’s post hoc test. \**P* < 0.05.

Table S1. Composition of reconstituted pB, imvB and smvB BA pools used for functional studies.

## Materials and Methods

### Mice

All mice were C57BL/6J background and maintained under specific pathogen-free conditions at Scripps Florida and Dartmouth College. C57BL/6J (B6, JAX cat. no. 000664) mice and *Il17a^tm1Bcgen^*/J (IL-17A-GFP, JAX cat. no. 018472) mice were purchased from the Jackson Laboratory. *Tnf^△ARE/+^* mice^5^ were provided by Fabio Cominelli (Case Western Reserve University) and bred as heterozygotes, cohoused with B6 littermates. *Slc10a2*^-/-^ mice^47^ were obtained from Paul Dawson (Emory University). *Slc10a2*-deficient *Tnf^△ARE/+^* (*Tnf^△ARE/+^*;*Slc10a2*^-/-^) mice were generated via crossing *Tnf^△ARE/+^* mice with *Slc10a2*^-/-^ mice. Sex- and age-matched mice were co-housed at weaning and analyzed at 16-weeks and 30-weeks of age. All experiments and procedures were approved by the Institutional Animal Care and Use Committees at Scripps Florida and Dartmouth College.

### Cholestyramine and tβMCA treatment

Rodent chow containing 2% cholestyramine (CME, Sigma-Aldrich) was made by Teklad. B6 and *Tnf^△ARE/+^* mice were co-housed at weaning with standard rodent chow until 25-weeks of age, followed by CME-supplemented chow for 5 weeks. For acute tβMCA treatment, B6 and *Tnf^△ARE/+^* mice were co-housed at weaning until 13-weeks of age, mice were treated with the 1^st^ dose of tβMCA at the concentrations of 0, 0.2, 1, and 10 mg/dose (dissolved in 100 μL PBS) by oral gavage, followed by the 2^nd^ dose of tβMCA after 24 hours (hr). For two-week tβMCA treatment, B6 and *Tnf^△ARE/+^* mice were co-housed at weaning until 13-weeks of age, mice were supplemented with tβMCA at the concentrations of 0, 0.2, 1, and 10 mg/dose (dissolved in 100 μL PBS) by oral gavage every other day for two weeks. All mice were fasted for 4 hr before euthanizing by CO_2_ narcosis.

### Histology

Swiss-rolled ileum (10 cm, from distal to proximal ileum) was fixed in 10% neutral buffered formalin and paraffin-embedded. Hematoxylin and eosin (H&E) staining were performed on 5-μm thick section by Pathology Shared Resource, Dartmouth Cancer Center. Histology was evaluated and scored blindly on the basis of the below criteria: crypt architecture (normal, 0; severe crypt distortion with loss of entire crypts, 3), inflammatory cell infiltration (normal, 0; dense inflammatory infiltrate, 3), muscle thickening (base of crypt sits on the muscularis mucosae, 0; marked thickening, 3), goblet cell depletion (absent, 0; present, 1) and crypt abscess (absent, 0; present, 1). The sum of the above was multiplied to generate a total histological score. For immunohistochemistry staining of Asbt/*Slc10a2*, 7-cm section of the distal ileum was dissected longitudinally and fixed in 10% neutral buffered formalin. 5-μm thick sections were processed for immunohistochemical detection of Asbt/*Slc10a2*. All slides were imaged using Keyence BZ-X800 microscope (Keyence). % area of Asbt/*Slc10a2* expression was evaluated by Fiji ImageJ v2.14.0/1.54f.

### Reconstitution of physiological BA pools

BAs were purchased from Avanti Polar Lipids, Cambridge Isotope Laboratories, Cayman Chemical, Med Chem Express and Sigma-Aldrich. BA pools were reconstituted using the mean values of BA concentrations in smvB of B6 (42 BA species, 311 μM) and *Tnf^ΔARE/^*^+^ (38 species, 88 μM) mice, imvB of B6 mice (49 BA species, 44 μM) and pB of B6 mice (51 BA species, 21 μM) as indicated in **Table. S3**. Two BA species including allocholic acid 3-SO4 (allo-CA 3-SO3) and deoxycholic acid 24-Gluc (DCA-24-Gluc) were excluded in BA pools given the commercial unavailability. BA species with the concentrations less than 0.00001 μM were also excluded in BA pools. For dropout and add-in experiments of tβMCA and tα/ωMCA, smvB BA pool of B6 was reconstituted without tβMCA (103.6 μM) or/and tα/ωMCA (26.1 μM and 14.3 μM, respectively); smvB BA pool of *Tnf^ΔARE/+^* was first reconstituted without tβMCA or/and tα/ωMCA, tβMCA (103.6 μM) or/and tα/ωMCA (26.1 μM and 14.3 μM, respectively) were then added individually. For dropout experiments of tβMCA, tDCA and tLCA, smvB BA pools of B6 were reconstituted without tβMCA (103.6 μM), tDCA (3.6 μM) or tLCA (0.19 μM). All BA pools were aliquoted and stored at -20 °C.

### Single cell suspensions of epithelial and lamina propria cells

Preparation of single cell suspensions were performed as previously described.^48^ Briefly, ileum (10 cm) and colon were removed, opened longitudinally and rinsed with ice-cold PBS. Peyer’s patches were removed from ileum. Dissected ileum and colon were cut into pieces (∼1 cm) and epithelial cells were dissociated by incubating in RPMI 1640 containing 1% FBS, 1 M MgCl_2_ and 0.5 M EDTA with shaking at 280 rpm for 30 min at 37 °C, dissociation of epithelial cells was conducted twice. The remaining tissues were digested in RPMI medium containing 20% FBS and 0.2 mg/mL Collagenase III (Thermo Fisher Scientific) with shaking at 280 for 30 min at 37 °C, digestion process was conducted twice. The epithelial cell fraction and lamina propria fraction were filtered through a 100-μm cell strainer and resuspended in 40% Percoll solution with centrifugation at 3,000 g for 12 min at room temperature (RT). Cells were then washed and resuspended in PBS containing 2% FBS.

### Flow cytometry

Single-cell suspensions were stained with anti-mouse CD16/CD32 antibody (Biolegend) for 10 min at 4 °C. Dead cells were excluded with Zombie NIR Fixable Viability Kit (Biolegend). Cells were then stained with surface conjugated antibodies for 20 min at 4 °C for myeloid cells analyses. The staining antibodies for flow cytometry were purchased from Biolegend: CD8a BV570 (53-6.7), NK1.1 BV650 (PK136), lineage markers: EpCAM BV605 (G8.8), F4/80 BV605 (BM8), CD11b BV605 (M1/70), CD19 BV605 (6D5) and Ly6G BV605 (1A8), IL-17A Alexa Fluor 700 (TC11-18H10.1), TCRβ APC (H57-597), CD19 PE-Cy7 (6D5), FceRIα APC-Cy7 (MAR-1), F4-80 BV785 (BM8), Ly6C BV711 (HK1.4) and EpCAM PerCp-Cy5.5 (G8.8); BD Biosciences: CD3e BUV395 (145-2C11), Ly6G BUV395 (1A8), CD11c BUV563 (N418), TCRγδ BUV615 (GL3), CD4 BUV661 (RM4-5), TCRβ BUV737 (H57-597), CD90.2 BUV805 (30-H12), T-bet BV421 (4B10), CD62L BV480 (MEL-14), CD8b BV750 (H35-17.2), CD127 BV711 (SB/199), CD44 APC-Cy7 (IM7), I-A/I-E APC/Fire 810 (M5/114.15.2), RORγt PE-CF594 (Q31-378), CD11b FITC (M1/70), SiglecF Alexa Fluor 647 (E50-2440) and STAT3 Pacific Blue (4/p-STAT3); Thermo Fisher Scientific: CD45 Alexa Fluor 532 (30-F11), Foxp3 eFluor450 (FJK-16s) and GATA3 PE-Cy5 (TWAJ). For transcriptional factors and cytokines staining, cells were first incubated in RPMI 1640 containing 10% FBS (Gibico), 10 nM phorbol myristate acetate (PMA), 1 μM ionomycin and 10 μg/mL brefeldin A at 37 °C for 4 hr. All obtained from Sigma-Aldrich. Cells then were fixed and permeabilized with FOXP3/Transcription Factor Staining Buffer Set following the manufacturer’s instructions (Thermo Fisher Scientific) after incubating with surface conjugated antibodies. Briefly, cells were incubated with FOXP3 Fixation/Permeabilization working solution overnight at 4 °C, then stained with intracellular markers in 1× permeabilization buffer for 1 hr at 4 °C. For phosphorylation of STAT3 *in vitro*, cells were fixed in pre-warmed 4% paraformaldehyde (PFA) at 37 °C for 10 min, followed by permeabilization with ice-cold Phosflow Perm Buffer III (BD) at 4 °C for 30 min. Flow cytometry data were acquired on a Cytex Aurora (Cytex) and a MACSQuant Analyzer 10 (Miltenyi). Data was analyzed using FlowJo Software (Tree Star, Biosciences).

### Human blood and tissue samples

Human blood samples and tissue biopsies were obtained from Miller School of Medicine at University of Miami following protocols approved by Institutional Review Boards (protocol number 20190548 for blood and tissue biopsies for BA measurements and nanostring; protocol number 20081100 for scRNA-Seq). A written informed consent was obtained from all subjects. For BAs study, human blood samples were obtained from 17 pairs (dyads) of adult healthy controls and ileal Crohn’s diseases patients living in the same household and consumed an identical (catered) diet for 8 weeks, serum samples were collected after centrifuging at 2,000 g for 10 min at RT and stored at -80 °C. For mRNA expression of *SLC10A2*, terminal ileal, ascending colonic and sigmoid colonic biopsies were obtained from patients with ileal Crohn’s disease (iCD) or Ulcerative colitis (UC) and stored in RNAlater (Invitrogen) at -80 °C. For scRNA-Seq of human epithelial and lamina propria cells, intestinal biopsies from the ileum and ascending colon were collected from 25 CD patients and 20 healthy individuals during a routine colonoscopy procedure. Samples were stored at 4 °C in HypoThermosol and then processed for single-cell suspensions. Briefly, biopsies were processed in 1 mM EDTA and 10 mM DTT to removed intestinal mucus and excess epithelial cells. Tissues were then mechanically dissociated and enzymatically digested using collagenase and DNase I with shaking at 37 °C for 45 minutes, followed by filtration through a 70-µm strainer. Cell viability of at least 85% for all samples were confirmed by trypan blue.

### *In vitro* CD4^+^ T cell cultures

CD4^+^CD25^-^CD44^-^ T cells were isolated from the spleen and peripheral lymph nodes of IL17a-GFP mice and magnetically enriched using EasySep Mouse CD4^+^ T Cell Isolation Kit (Stem Cell Technologies) with biotin-conjugated CD25 (3 mg/mL, Clone PC61) and CD44 (3 mg/mL, Clone IM7) purchased from BioLegend. Cells were seeded (at 5 × 10^5^ cells/mL) in a 96-well round-bottom plate pre-coated with goat anti-hamster IgG (50 μg/mL, Invitrogen) in T cell medium (Advanced DMEM, 2 mM L-glutamine, 50 mM 2-mercaptoethanol, 10 mM HEPES, 100 U/ml penicillin and 100 mg/ml streptomycin) supplemented with 1 μg/mL anti-CD3e (Clone 145-2C11) and 0.25 μg/mL anti-CD28 (Clone 37.51). For Th0 culture, T cells were cultured with T cell medium without cytokines. For Th1 cell differentiation, T cells were cultured with the addition of 5 ng/mL IL-12 and 5 μg/mL anti-IL-4 (Clone 11B11). For Th2 cell differentiation, T cells were cultured with the addition of 10 ng/mL IL-4 and 5 μg/mL anti-IFNγ (Clone XMG1.2). For non-pathogenic Th17 (npTh17) cell differentiation, T cells were cultured with the addition of 30 ng/mL IL-6 and 3 ng/mL TGFβ. For pathogenic Th17 (pTh17) cell differentiation, T cells were cultured with the addition of 30 ng/mL IL-6, 1 ng/mL IL-1β and 10 ng/mL IL-23. For inducible Tregs (iTregs) differentiation, T cells were cultured with the addition of 5 ng/mL TGFβ and 2 ng/mL IL-2. For smvB BA pools of B6 and *Tnf*^△ARE/+^ treatments, cells were treated with BA pools on day 0, IL17a-GFP^+^ cells and mRNA expression of transcriptional factors and cytokines of CD4^+^ T cells were analyzed on day 3. For BAs drop out experiments, npTh17 cells were treated with BA pools on day 0 and IL-17A-GFP^+^ cells were analyzed on day3. For proliferation assay, cells were labelled with CellTrace Violet (Invitrogen), and division index was analyzed following the manufacturer’s instructions. For phosphorylation of STAT3, CD4^+^CD25^-^CD44^-^ T cells were treated with BA pools on day 0 and MFI of pSTAT3 was analyzed at 0 hr, 0.5 hr, 2 hr, 6 hr, 12 hr and 24 hr. For kinetics of RORγ expression and IL-17A-GFP^+^ cells, npTh17 cells were treated with BA pools on day 0, RORγt expression was analyzed at 6 hr, 24 hr, 48 hr, 66 hr and 72 hr, IL-17A-GFP^+^ cells were analyzed at 48 hr, 60 hr, 66 hr and 72 hr.

### scRNA-Seq

#### Cell isolations

Human epithelial and lamina propria cells from Crohn’s diseases samples were generated as described above. Single-cell suspensions of ileal epithelial cells from co-housed B6, *Slc10a2*^-/-^, *Tnf^△ARE/+^* and *Tnf^△ARE/+^*;*Slc10a2*^-/-^ mice at 15 weeks old were processes as described above. 2 mice were pooled per sample. Dead cells were excluded via Dead Cell Removal Kit (Miltenyi Biotec) according to the manufacturer’s instructions. Briefly, cells were incubated in Dead Cell Removal MicroBeads for 15 min at RT, then processed to magnetic separation with LS columns. Flow-through containing live cells was collected and viability of at least 90% for all samples were confirmed by trypan blue. Single-cell suspensions of ileal lamina propria cells from co-housed B6, *Slc10a2*^-/-^, *Tnf^△ARE/+^* and *Tnf^△ARE/+^*;*Slc10a2*^-/-^ mice at 15-weeks of age were performed as described above. Cells were incubated with anti-mouse CD16/32 (BD, Clone 2.4G2) for 10 min at 4 °C, followed by TotalSeq-C Hashtag antibodies (BioLegend) for 30 min at 4 °C. CD45^+^TCRβ^+^CD4^+^ cells (CD4^+^ T cells) and CD45^+^CD4^+^ cells (non-CD4^+^ T cells) were then sorted into 5 μL PBS containing 10% FBS in an Eppendorf tube using FACSAria Fusion Cell Sorter (BD). Viability of at least 80% for all samples were confirmed by trypan blue.

#### Processing of raw scRNA-Seq data

Probe-based Amplified RNA Single-cell Expression (PARSE) profiling was applied for human scRNA-Seq. Briefly, single-cell suspensions were fixed and permeabilized using the Parse EvercodeTM Fixation kit (Parse), then stored at -80°C until barcoding. Transcript barcoding and library preparation were performed using the Parse EvercodeTM WT (Whole Transcriptome) Mega kit. PARSE libraries were sequenced on Illumina NovaSeq X Plus to a depth sufficient to achieve robust detection of cellular transcripts. Raw sequencing reads were processed using the Parse Biosciences cloud-based pipeline hosted on Amazon Web Services (AWS). The pipeline performs barcode decoding for split–pool combinatorial indexing, read alignment to a reference genome, unique molecular identifier (UMI) collapsing, and gene-level quantification. Cell barcodes were identified and filtered to exclude low-quality or background-associated barcodes. Digital gene expression (DGE) matrices were generated for each sample, and the filtered matrices located in the all-sample/DGE filtered directory were used for downstream single-cell analyses using CLC Genomic Workbench.

Mice scRNA-seq libraries were prepared using the Chromium Single Cell 5’ Library Kit (10X Genomics) at Single Cell Genomics Core, Dartmouth Cancer for Quantitative Biology. On-chip multiplexing (OCM) was applied to scRNA-Seq on ileal epithelial fraction in mice. Libraries were sequenced on an Illumina NextSeq 2000. CellRanger v8.0.0 (mice lamina propria cells) and v9.0.0 (mice epithelial cells) were used to perform quality control, sample demultiplexing, barcode processing, alignment and single cell 5’ gene counting. The Raw reads were mapped to the mm10 mouse reference genome. Filtered barcode matrices were quantified and used to build the gene expression matrix.

#### Pre-processing, doublets removal, and quality filtering

AnnData objects of mice epithelial cells and human epithelial and lamina propria cells, and Seurat object of mice lamina propria cells were created and further processed using python v3.12.2 Scanpy v.1.10.3 and R v4.5.1 Seurat package v5.3.1, respectively. Cells that expressed less than 100 genes and genes that were not detected in at least three (human) or ten (mice) single cells were filtered out. Low-quality and dying cells were removed via assessing nFeature RNA, nCount RNA, and percentage of mitochondrial gene expression (% MT in human, % mt in mice). Cells with 300 < nFeature RNA < 5,000, 300 < nCounts < 15,000, and % MT < 15 in human epithelial and lamina propria cells, 800 < nFeature RNA < 7,500, 200 < nCounts < 20,000, and % mt < 10 in mice epithelial cells, and 200 < nFeature RNA < 6,000, 600 < nCounts < 30,000, and % mt < 10 in mice epithelial cells were retained. Doublets were detected and filtered out using Scrublet v0.2.3 (Scanpy) and scDblFinder v1.24.0 (Seurat), respectively.

#### Analysis of cell clusters and differentially expressed genes

The filtered data further analyzed following the clustering tutorial of Seurat (https://satijalab.org/seurat/) and Scanpy (https://scanpy.readthedocs.io/en/stable/index.html). Briefly, the filtered cells were log-normalized and scaled (LogNormalize in Seurat, pp.normalize_total in Scanpy), followed by principal component analysis (PCA) on highly variable features (SCTransform in Seurat, pp.highly_variable_genes in Scanpy). Human scRNA-Seq data was further integrated using pp.harmony_integrate function in Scanpy. Nearest neighbor graphs were constructed using the FindNeighbors function in Seurat or sc.pp.neighbors in Scanpy with the numbers of significant principal components identified from PCA analysis. Clusters were identified using FindClusters function in Seurat with resolution of 0.6 for mice lamina propria cells, 0.80 for mice epithelial cells, and 0.05 for human epithelial and lamina propria cells. Sub-clustering of IECs in human scRNA-Seq and absorptive IECs in mice scRNA-Seq were further analyzed. Briefly, IECs were extracted and re-analyzed from normalization, highly variable genes selection, PCA, and clustering analysis with the resolution of 1.20 and 0.80, respectively. Cell annotations and cluster names were performed by the marker genes.^20,49–52^ Clusters were visualized with UMAP using the Seurat packages or Scanpy. Differentially expressed genes were performed using Wilcoxon rank-sum test by FindMarker function in Seurat or tl.rank_genes_group in Scanpy. Significantly upregulated and downregulated genes were defined as adjusted *P* value (*P*adj) < 0.05 with the threshold of 0.25 or 0.50 for log_2_fold change. All visualization were performed using R packages Seurat or Scanpy.

### Liquid Chromatography with tandem mass spectrometry (LC-MS/MS)

Samples including siLC, smvB, feces and pB were collected, weighed and stored at -80 °C before BAs quantification. BAs analysis was performed by Creative Proteomics as previously described ^4^. Briefly, for blood, 20 µL samples were mixed with 80 µL internal standard solution (50% methanol/50% water containing 0.01% formic acid and deuterated BAs) and 900 µl water. Samples were loaded onto a polymeric reversed-phase SPE cartridge after sonication. BAs were eluted with 1 mL methanol, dried under nitrogen gas flow, and reconstituted in 80 µL of 50% acetonitrile. For siLC and feces, samples were homogenized in 20 µL of 70% acetonitrile per 1 mg tissue using a Mixer mill MM 400 (Retsch, Haan, Germany) at 30 Hz for 3 min and centrifuged; the supernatant was diluted with internal standard solution containing all the target BAs. BAs measurement was performed on an Agilent 1290 UHPLC system (Agilent, CA, USA) coupled to a Sciex 4000 QTRAP mass spectrometer (Sciex, Marlborough, MA, USA). tβMCA quantification in smvB and cecum was performed by Analytical Chemistry Core at Harvard Medical School. LC-MS was operated in negative electrospray ionization mode. The concentration of tβMCA were calculated using a standard curve.

### NR reporter assays

Multiplex NR reporter assays were performed as fee-for-service, at Attagene, Inc., as previously reported.^53^ Briefly, each human NR ligand-binding domain (LBD) fused with the SV40 promoter and GAL4 DNA-binding domain (DBD), was expressed in a Chinese hamster ovary (CHO) cell line (ATCC, Manassas, VA, USA) transfected with human ASBT/*SLC10A2*. The GAL4-NR vector includes a GAL4 reporter transcription unit (RTU) containing an identical reporter sequence with an HpaI restriction enzyme recognition site at a different position under the control of GAL4 DBD binding promoter. A GAL4-expressiong vector lacking the NR LBD and a TATA module without an expression vector were used as negative and internal controls, respectively. After separate transfections, cells were pooled and cultured in DMEM medium (Gibco) supplemented with 1% charcoal-treated FBS (Hyclone, Logan, UT, USA) in the presence of equivalent concentrations of smvB BA pools of B6 or *Tnf^△ARE/+^* mice or tCDCA for 24 hr. Total RNA was isolated with the PureLink Pro 96 total RNA Purification Kit (Invitrogen) and then reverse-transcribed into cDNA using RT-PCR, labeled with 6-carboxyfluorescein (6-FAM) using a 6-FAM-labeled RTU-specific primer, digested with HpaI, and separated by capillary electrophoresis (CE) using a 3130xl Genetic Analyzer (Applied Biosystems, Waltham, MA, USA). CE signals were normalized to TATA module signals and NR ligand activities were calculated relative to vehicle (DMSO)-treated cells.

### NR pulldown studies

The pulldown experiments were performed as previously described.^54^ Briefly, the ligand-binding domain (LBD) of human FXR or RORγ containing a TEV-cleavable 6× histidine-tag (his-tag) were expressed in *Escherichia coli* BL21 (DE3). Bacterial cell lysates were incubated with reconstituted smvB BA pools of B6 or *Tnf^△ARE/+^* mice in the presence of HIS-select nickel magnetic agarose beads (Sigma-Aldrich) for 1 hr at 4 °C, followed by 30 min at RT. His-tagged LBD-ligand complexes were harvested using a magnetic stand and analyzed by LC-MS. Samples were first loaded onto a reversed-phase column (Accucore Vanquish C18+, 1.5 μm, 50 × 2.1 mm, Thermo Scientific) using a UPLC system in methanol and eluted with a gradient starting at 90% mobile phase A (H_2_O with 0.1% formic acid) and 10% mobile phase B (methanol with 0.1% formic acid) at a flow rate of 0.25 ml/min. The gradient was increased to 80% mobile phase B at 2 min, 85% mobile phase B at 7 min and 100% mobile phase B at 8 min, then held isocratic for 1 min before returning to 90% mobile phase A over 0.5 min, followed by an additional 2.5 min of isocratic flow. The column and sample compartments were maintained at 40 °C and 10 °C, respectively. Data acquisition was performed in full MS and parallel reaction monitoring (PRM) mode. Full MS scans were recorded at a resolution of R = 70,000, with a maximum ion count of 3 × 10^6^ collected within 200 ms. Metabolomic data supporting the findings of this study have been deposited in MassIVE under accession ID: MSV000100375.

### Metagenomics

Cecal metagenomics were performed by TransnetYX (Cordova). Cecal DNA was extracted using Qiagen DNeasy 96 PowerSoil Pro QIAcube HT kits (Qiagen). Sequencing libraries were prepared with Watchmaker DNA Library Prep kits with Fragmentation (Watchmaker) and sequenced on an Illumina NovaSeq instrument (Illumina) using shotgun sequencing at a depth of 2 million paired end reads (2 × 150 bp). Acquired data were analyzed using the One Codex database. Metagenomic reads were processed with atlas v.2.13.0.^55^ Briefly, reads were filtered for mouse host contamination using STAR suite v.2.7.2b,^56^ error-corrected and merged before assembly with metaSpades v.3.15.3.^57^ Contigs were binned using maxbin2 v.2.2 and predicted MAGs were clustered at 95% average nucleotide identity, yielding 34 representative genomes. Gene prediction for each genome was performed using eggNOG v.2.1.^58^ α-diversity and β-diversity of metagenomic communities were calculated based on MAG relative abundance outputs from the atlas workflow using the vegan R package v.2.6-4.

### Nanostring

Total RNA from human biopsies and mouse liver and terminal ileum was isolated using TRIzol (Invitrogen) or RNeasy Mini Kits (Qiagen) following the manufacturer’s instructions. RNA samples were run on nCounter platforms using custom codesets (Nanostring) by Dartmouth Cancer Center, Genomics Shared Resource. Raw data was normalized using 3 housekeeping genes (*Rps19*, *Gapdh*, and *Actb*) and analyzed using nSolver software (Nanostring).

### qPCR

SFB colonization levels were performed by qPCR as previously described.^59^ Briefly, bacterial DNA was extracted from fecal pellets with DNA extraction buffer (200 mM Tris, 200 mM NaCl, 20 mM EDTA), 20% SDS, and phenol:chloroform:isoamyl alcohol (25:24:1). DNA-containing aqueous phase is collected and subjected to two additional phenol:chloroform:isoamyl alcohol (25:24:1) extraction to further remove organic contaminants. RNA was removed using RNase A (Thermo Fisher). SFB-specific primers Primer sequences were as follows: SFB, 5’-GACGCTGAGGCATGAGAGCAT-3’, 5’-GACGGCACGGATTGTTATTCA-3’; Eubacteria (UNI), 5’-ACTCCTACGGGAGGCAGCAGT-3’, 5’-ATTACCGCGGCTGCTGCG-3’. SFB abundance was normalized to total bacterial levels as determined by the UNI primers. Positive control was obtained from fecal pellets of mice colonized with SFB. For mRNA expression of transcriptional factors and cytokines from CD4^+^ T cells *in vitro*, RNA was extracted using Trizol and cDNA was synthesized using a High-Capacity cDNA Reverse Transcription Kit (Life Technologies). Probes for mouse genes included: *Actb* (Mm00607939_s1), *Rorc* (Mm01261022_m1), *Il17a* (Mm00439619_m1), *Tbx21* (Mm00450960_m1), *Ifng* (Mm01168134_m1), *Gata3* (Mm00484683_m1), *Il4* (Mm00445259_m1), *Foxp3* (Mm00475162_m1) and *Il10* (Mm01288386_m1). qPCR was performed on a QuantStudio 6 Pro (Applied Biosystems).

### Protein expression for TR-FRET assays

DNA encoding the human RORγ LBD (UniProt ID P51449; NR1F3; residues 260–518) was synthesized by TwistBioscience into pET21 expression vector as a 3C cleavable N-terminal 6x-polyhistidine tag fusion protein. Recombinant 6xHis-hRORγ LBD was expressed in *Escherichia coli* BL21(DE3) cells. Cells were inoculated with the overnight preculture and grown for 5 hours at 37 °C, then 1 hr at 30 °C and then 16 hr at 18 °C. Cells were harvested by centrifugation at 4,000 rpm at 4 °C for 30 minutes and washed with PBS followed by centrifugation at 4,000 rpm at 4 °C for 30 minutes. Cells were then resuspended in lysis buffer (40 mM Phosphate buffer pH 7.4, 500 mM KCl, 10% glycerol, 0.05% CHAPS and 15 mM Imidazole) supplemented with 10 µg/mL DNAse I, 10 µg/mL Lysozyme, 10 µg/mL Pepstatin A, 10 µg/mL Leupeptin, 1 mM PMSF and 2 mM TCEP and lyzed by sonication during 5 minutes, 75% amplitude with 15 seconds on, 25 seconds off pulses. The lysate was then cleared by centrifugation for 35 minutes at 15,000 rpm at 4 °C and filtered with a 0.22 µm filter. To the soluble fraction, 5 mL of NiNTA slurry was added and kept at 4 °C in a rotator for 1 hr for protein binding to the beads. Beads were then washed 3 times with lysis buffer and eluted with lysis buffer supplemented with 485 mM imidazole. Protein was dialyzed to a buffer containing 20 mM HEPES pH 7.4, 150 mM NaCl and 0.5 mM EDTA overnight. Protein purity was checked on a 12% SDS-PAGE gel prior to use for biochemical experiments.

### TR-FRET co-regulator interaction assays

Time-resolved fluorescence resonance energy transfer (TR-FRET) coregulator peptide interaction assays were performed in low-volume black 384-well plates (Greiner) using a 20 μL final well volume. Each well contained 4 nM protein (6xHis-hRORγ LBD), 1 nM LanthaScreen Elite Tb-anti-His Antibody (ThermoFisher), and 400 nM FITC-labeled co-regulator peptides in a buffer containing 20 mM HEPES pH 7.4, 150 mM NaCl, and 0.5 mM EDTA, 1 mM TCEP and 0.005% Tween 20 (FRET buffer). Ligand stocks were prepared via serial dilution in DMSO, then diluted 1:100 in the FRET buffer, added to wells in duplicate (90 μM highest final ligand concentration), and plates were read using BioTek Synergy Neo multimode plate reader after incubation for 1 hr at 25 °C. The Tb donor was excited at 340 nm; its emission was measured at 495 nm and the acceptor FITC emission was measured at 520 nm. Data were plotted using GraphPad Prism as TR-FRET ratio (520 nm/495 nm). *vs.* ligand concentration and fit to a sigmoidal dose–response equation.

### Thermal shift assays

The thermostabilities of RORγ-LBD in the presence of DCA, LCA, tDCA, or tLCA were assayed using a Thermofluor-type assay. Briefly, 2 μL of 30 μM purified wild-type RORγ-LBD, was dispensed into 96-well plates (∼3 μM protein per well). Compounds in 100% (v/v) DMSO were serial diluted with DMSO and added into wells (1 μL for each reaction; final concentration of DMSO 1% (v/v)). 15 μL of thermal shift assay buffer was added into wells to make 18 μL of reaction solutions. Proteins and ligands were then incubated at 4°C for 2 hr. A 2 μL aliquot of 100-fold SYPRO Orange dye (Sigma-Aldrich) was added into each well and then wells were sealed. Dye and protein solutions were mixed by gently rotating for 15 s and spun down at 1,500 × g for 2 min. Protein melting curves were measured using the QuantStudio 6 Flex Real-Time PCR System (Thermo fisher) with the following program: 2 min at 25 °C, ramp to 95 °C at 0.3 °C•s^-1^, 2 min at 95 °C, excitation and emission wavelengths 483 nm and 568 nm, respectively. The melting temperatures (T_m_s) of the RORγ-LBD protein in the absence of ligands provided the baseline T_m_ (T_m_0).

### Flexible side chain molecular docking

Bile acid ligands were prepared for molecular docking using MGL AutoDockTools (ADT).^60^ Gasteiger partial charges were assigned, non-polar hydrogens were merged, and rotatable bonds were defined for each ligand prior to conversion to PDBQT format. An AlphaFold-predicted model of the human RORγ ligand-binding domain (UniProt ID P51449; NR1F3; residues 265–518) was used as the receptor. The receptor structure was prepared for docking in ADT by adding polar hydrogens, assigning Gasteiger charges, merging non-polar hydrogens, and assigning AutoDock atom types. Missing side-chain atoms were rebuilt where necessary prior to docking. To account for local side-chain flexibility within the ligand-binding pocket, selected residues were treated as flexible during docking, including Y281, L287, H323, L324, Y330, R364, M365, R367, F377, F378, F388 and L391. The docking grid box was centered on the orthosteric ligand-binding site defined by the digoxin-bound RORγ LBD crystal structure (PDB ID: 3B0W)^31^ with dimensions of 50Å x 42Å x 36Å. Docking simulations were performed using AutoDock-Vina version 1.2.7^61,62^ in batch mode with both rigid-receptor and flexible-residue protocols for the previously defined residues. For each ligand, 20 binding poses were generated using an exhaustiveness value of 32. The resulting poses were ranked by predicted binding affinity and visually inspected for consistency with known RORγ ligand binding modes. Predicted binding energies from AutoDock Vina were used for relative ranking and not interpreted as absolute free energies of binding. Structures were analyzed and figures prepared The PyMOL Molecular Graphics System version 3.0 (Schrödinger, LLC).

### Statistical analyses

All statistical analysis tests were performed using Prism v10.6.0 (GraphPad). *P* values were determined using Wilcoxon matched-pairs signed rank test, paired *t*-tests, unpaired *t*-tests, Brown-Forsythe and Welch ANOVA with Dunnett’s T3 multiple comparisons test, one-way ANOVA followed by Tukey’s multiple comparison test or Šídák’s post hoc test, two-way ANOVA analysis followed by Tukey’s multiple comparison test and Wilcoxon rank-sum test as indicated in the figure legends. BAs analyses were performed on R and figures were generated using R packages ggplot2^63^ and circlize.^64^ PCA was performed using singular value decomposition. BA variables were centered and scaled to mean of 0 and variance of 1. Effect of BA on principal components was determined by extracting the eigenvectors of the PCA. Significant differences were calculated using the paired Wilcox signed-rank and unpaired rank-sum tests with false discovery rate (*P*_adj_) calculation.

### Resource availability

#### Lead contact

Additional correspondence and requests for materials should be addressed to the lead contact, Mark S. Sundrud.

#### Materials availability

All materials generated in this paper will be made available by the lead contact upon request.

#### Data and code availability

All data are present in the paper or supplementary materials. Mouse scRNA-seq data reported here are available at the National Center for Biotechnology Information (NCBI) under accession codes GSE316588 and GSE316586. Human scRNA-seq data are accessible upon request. This paper does not report new code. Any additional information required to reanalyze the data reported in this paper is available from the lead contact upon request.

