## Supplemental Table 1 for "Ileitis abolishes tolerogenic functions of the enterohepatic bile acid pool"

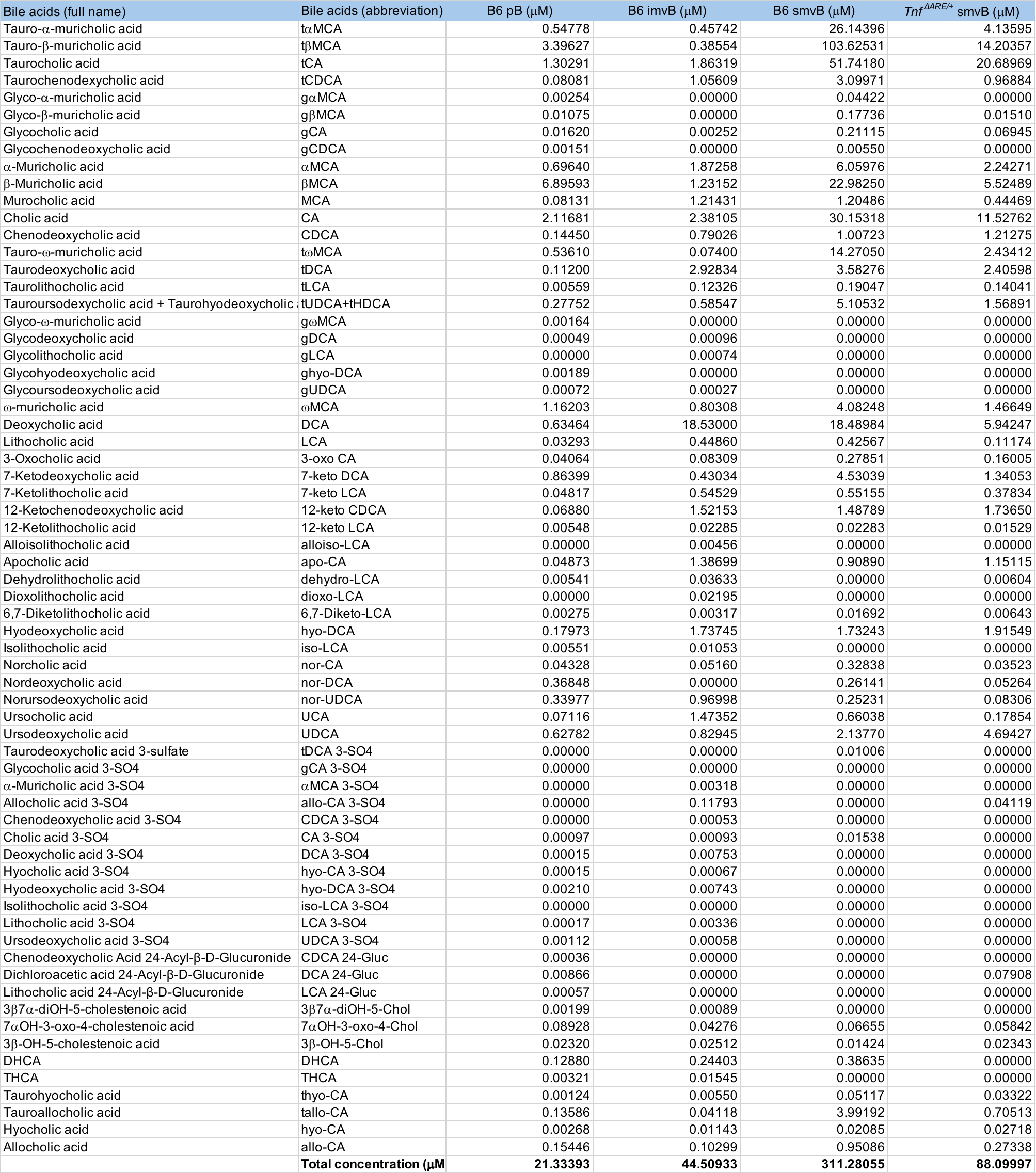
**Table S1.** **Composition of reconstituted pB, imvB and smvB BA pools used for functional studies.** Note that values listed as '0' include concentration < 0.00001 μM.
